# Exploration of novel αβ-protein folds through *de novo* design

**DOI:** 10.1101/2021.08.06.455475

**Authors:** Shintaro Minami, Naohiro Kobayashi, Toshihiko Sugiki, Toshio Nagashima, Toshimichi Fujiwara, Rie Koga, George Chikenji, Nobuyasu Koga

**Affiliations:** Protein Design Group, Exploratory Research Center on Life and Living Systems (ExCELLS), National Institutes of Natural Sciences (NINS), Okazaki, Aichi 444-8585, Japan; Institute for Protein Research, Osaka University, Suita, Osaka 565-0871, Japan; RIKEN, Yokohama, Kanagawa 230-0045, Japan; Department of Applied Physics, Graduate School of Engineering, Nagoya University, Nagoya, Aichi, 464-8603, Japan; SOKENDAI, The Graduate University for Advanced Studies, Shonan Village, Hayama, Kanagawa 240-0193, Japan; Research Center of Integrative Molecular Systems, Institute for Molecular Science (IMS), National Institutes of Natural Sciences (NINS), Okazaki, Aichi 444-8585, Japan

## Abstract

Most naturally occurring protein folds have likely been discovered^1–3^. The question is whether natural evolution has exhaustively sampled almost all possible protein folds^4^, or whether a large fraction of the possible folds remains unexplored^5–7^. To address this question, we introduce a set of rules for β-sheet topology to predict novel folds, and carry out the systematic de novo protein design for the novel folds predicted by the rules. The rules predicted eight novel αβ-folds with a four-stranded β-sheet, including a knot-forming one. We designed proteins for all the predicted αβ-folds and found that all the designs are monomeric with high thermal stability and fold into the structures close to the design models, demonstrating the ability of the set of rules to predict novel αβ-folds. The rules also predicted about twelve thousand novel αβ-folds with five- to eight-stranded β-sheets; the number is far exceeding the number of αβ-folds observed so far. This result suggests that the enormous number of αβ-folds are possible but have not emerged or become extinct due to evolutionary bias. The predicted novel folds should open up the possibility of designing functional proteins of our interests.

---

The structural diversity of proteins underlies their functional variety. The overall protein structure is determined by “fold”, the spatial arrangement of and connections between secondary structure elements. The number of naturally occurring protein structures solved and deposited in the Protein Data Bank (PDB) is currently more than hundreds of thousands and still continues to grow. On the other hand, the discovery of novel protein folds recently has become a rare event^1–3^, suggesting that almost all folds existing in nature have already been found. However, this does not necessarily indicate that we have uncovered all folds accessible to the polypeptide chain. Although debated^4–7^, natural evolution may only have sampled a small fraction of the possible fold space: there possibly exists a vast fold space not explored by natural evolution^5–7^.

We investigate the possibility through de novo protein design for the folds that have not been sampled by natural evolution. Recently developed principles for designing protein structures have made possible the design of a wide range of new proteins from scratch^9–14^, allowing us to explore the huge sequence space beyond evolution. However, in terms of the fold space, the exploration has been limited to naturally occurring protein folds^9–14^ except for one new fold of a protein called Top7^8^. To explore the fold space beyond evolution, a “map” to search for the folds that are possible but not observed in nature (i.e., novel folds) is indispensable; we derive a set of rules for β-sheet topology to predict novel folds. Here, we carry out systematic exploration of novel αβ-folds through *de novo* protein design, guided by the rules.

## αβ-folds not observed in nature

The αβ-folds, most of which are involved in enzymatic functions^18^, account for more than half of the currently identified protein folds^19^. We first sought to identify unobserved αβ-folds with a three- to eight-stranded open β-sheet (i.e., a β-sheet that does not form a barrel). We defined αβ-folds in a more abstract way than the original fold definition based on their β-sheet topology: the number of constituent β-strands in a β-sheet, and their order along the linear chain and orientations (Fig. 1a). Moreover, we considered only the folds with the right-handed strand connections between parallelly aligned β-strands as per Richardson’s rule^15^ (Fig. 1b). These lead to *n*! × 2*^n^*^-2^ patterns in total for αβ-folds for an *n*-stranded β-sheet, resulting in the finding of an enormous number of αβ-folds not observed in nature (Fig. 1c). However, apparently, all of the identified unobserved folds are not possible. For example, the fold shown at the bottom in Fig. 1a would not be possible because the two β-strand connections are overlapped. Therefore, we introduced a criterion that predicts possible αβ-folds among all patterns of the β-sheet topologies based on a set of rules for β-sheet topology.

**Fig. 1.**
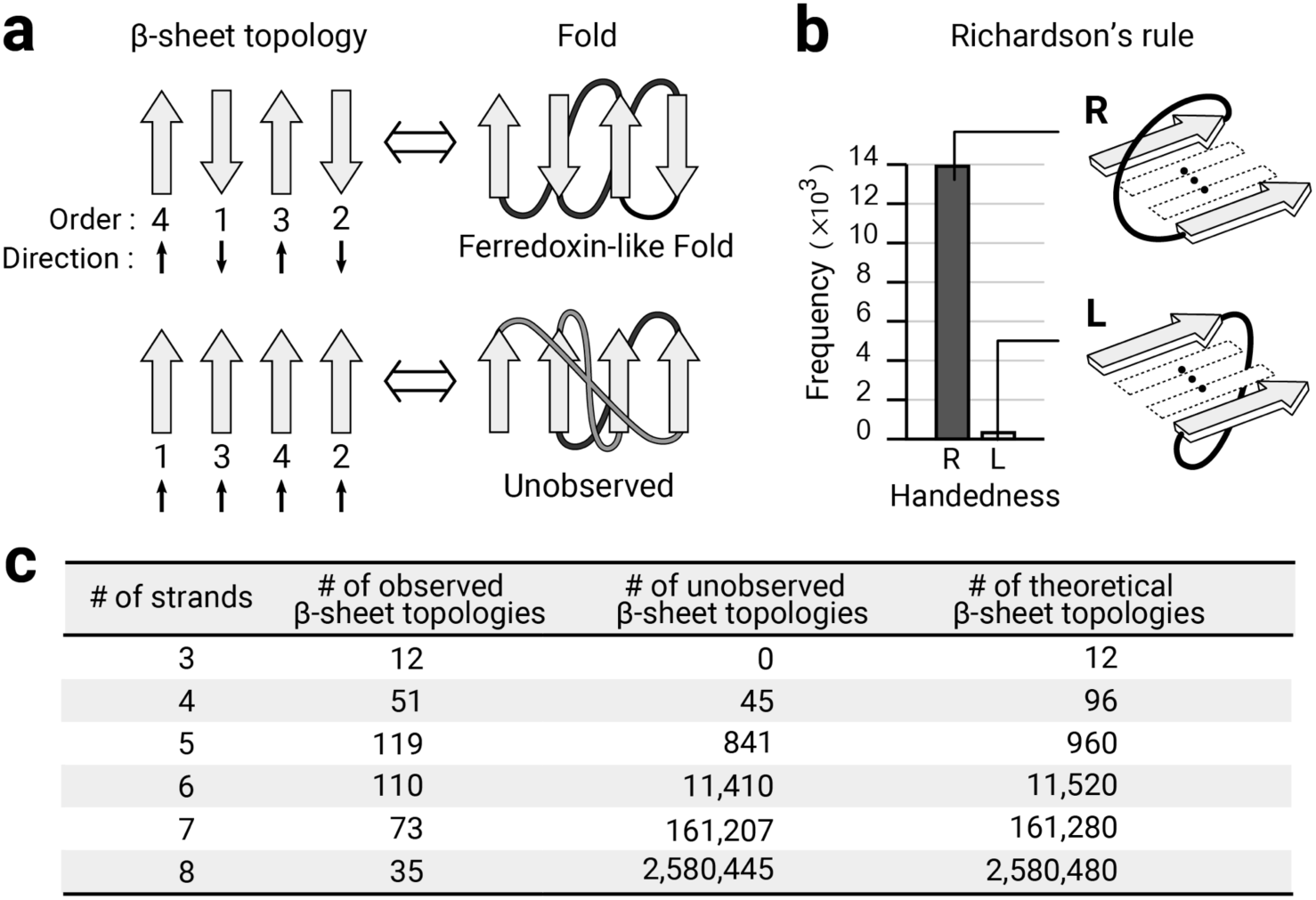
Observed and unobserved β-sheet topologies in nature. **a**, αβ-folds defined based on the β-sheet topology (the number of constituent β-strands, and their order and orientations) and Richardson’s right-handed strand connections^15^ shown in (**b)**. Two examples for a four-stranded β-sheet are illustrated: (upper panel) a frequently observed β-sheet topology in nature, and its corresponding fold, Ferredoxin-like fold; and (lower panel) an unobserved β-sheet topology in nature, and its corresponding fold. Each β-strand is numbered by the secondary structure order along the linear chain. The β-strand connections coloured in gray are on the front side of the β-sheet and those in black are behind. **b**, Richardson’s rule on the connection handedness of para-β-X-β motifs^15^. The right-handed strand connection (dark gray bar) is dominantly observed in naturally occurring proteins, compared to the left-handed one (light gray bar). **c**, The table shows the number of observed, unobserved, and theoretically possible β-sheet topologies for each number of constituent β-strands in a β-sheet.

## Rules for β-sheet topology

The set of rules described below are derived from conformational preferences of β-X-β motifs in naturally occurring protein structures, where X represents any backbone conformation (see Methods): the connection jump-distance rule for single β-X-β motifs, and the connection overlap and connection ending rules for pairs of β-X-β motifs.

### Connection jump-distance rule

The number of intervening β-strands between the two β-strands for para-β-X-β motifs (i.e., jump-distance) is less than four, and that for anti-β-X-β motifs is less than two (Fig. 2a); an exception is the anti-β-X-β motifs with two intervening β-strands, contained in the Greek-key β-sheet topology and its circular permutations (a dotted bar in Fig. 2a and the topologies with asterisks in Fig. 3c).

**Fig. 2.**
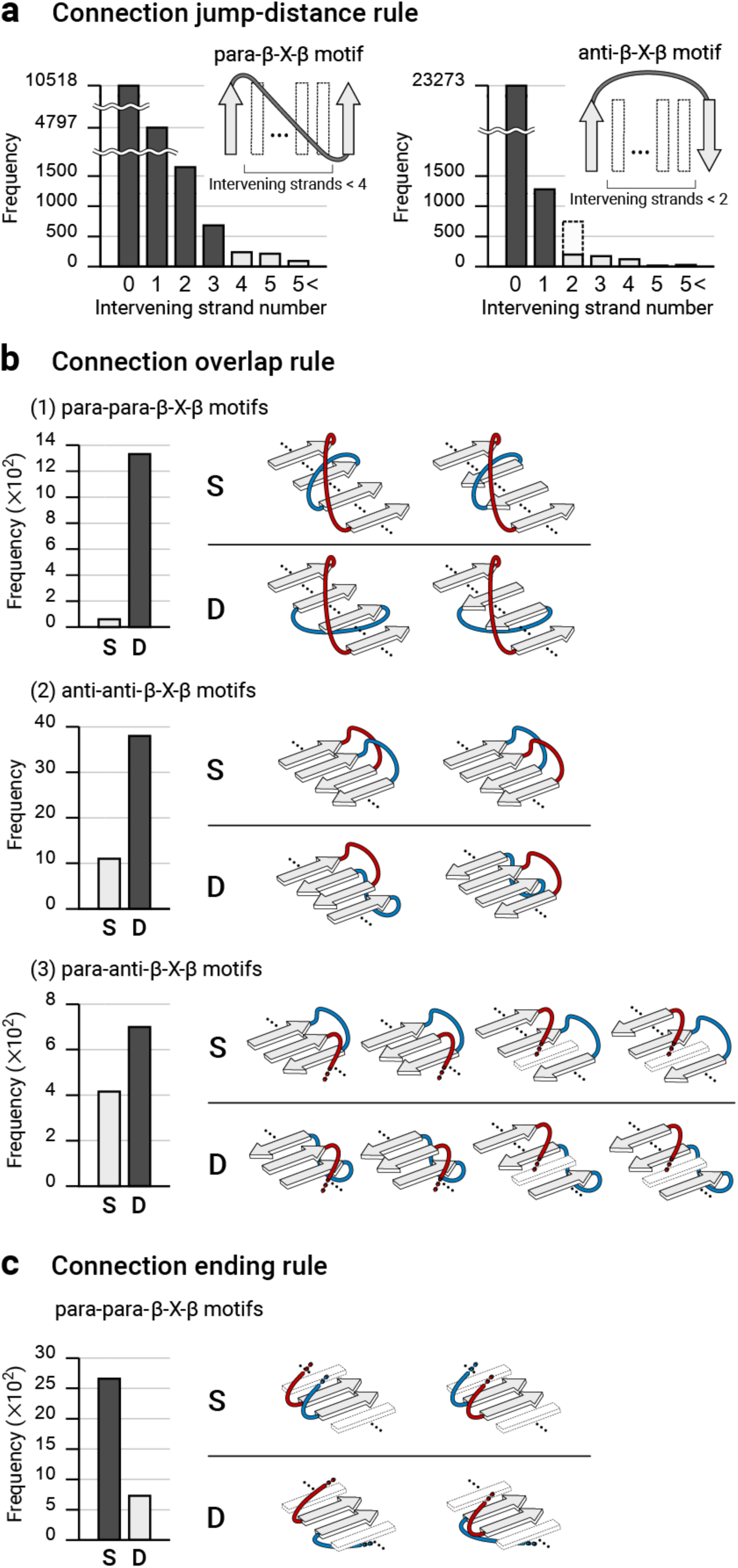
Rules for β-sheet topology. **a**, Connection jump-distance rule. Para-β-X-β and anti-β-X-β motifs are illustrated. The number of intervening strands between the two β-strands in para-β-X-β motifs is mostly less than four, and that for anti-β-X-β motifs is less than two; an exception is the anti-β-X-β motifs with the number of two, included in the Greek-key β-sheet topology and its circular permutations (the dotted box in the right histogram) (See topologies with an asterisk in Fig. 3d). . The same preferences are reported in the previous study^16^. We revisited them using the current PDB. **b**, Connection overlap rule. The preferred and not-preferred β-sheet topologies for three types for pairs of β-X-β motifs (para-para-, anti-anti-, and para-anti-β-X-β motifs) are illustrated. The D-type β-sheet topologies (loops are located on different sides of each other) are more frequently observed than the S-type ones (loops on the same side). Similar rules have been known for para-para-β-X-β motifs^62^. For anti-anti-β-X-β motifs, there has been a rule called “pretzels”^21^, but this rule prohibits both S- and D-types without the distinction. **c**, Connection ending rule. The two types of β-sheet topologies for pairs of para-β-X-β motifs are illustrated, in which the second strands of the two motifs are adjacent and parallelly aligned. The S-type β-sheet topologies are more preferred than the D-type topologies.

**Fig. 3.**
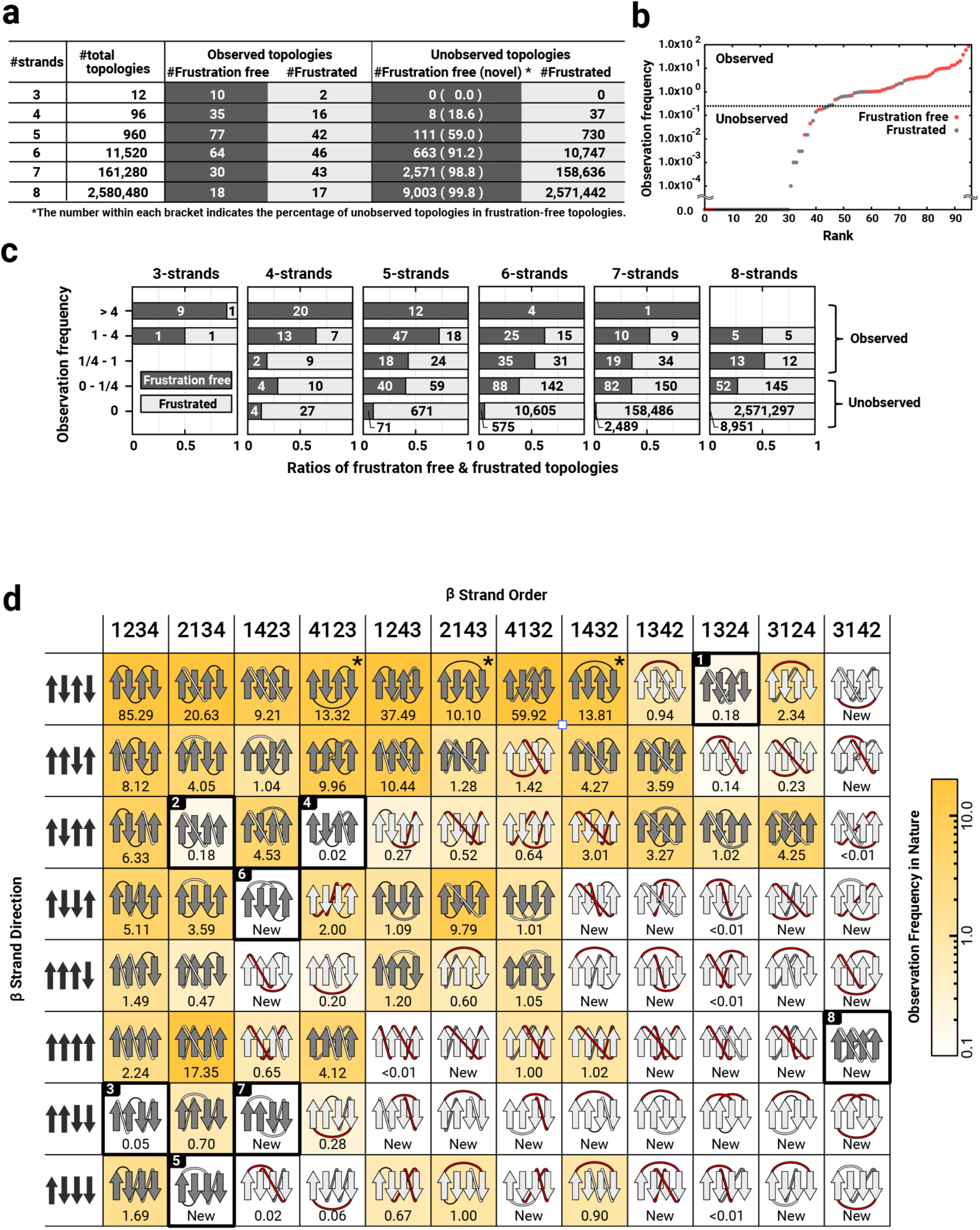
Distributions of frustration-free and frustrated β-sheet topologies in nature. **a**, The number of frustration-free and frustrated β-sheet topologies in each observed and unobserved topology in nature, for each number of constituent β-strands in a β-sheet. **b**, Observation frequencies of all possible 96 topologies for 4-stranded β-sheets, sorted in order of each observed frequency. Red points represent frustration-free topologies and grey points, frustrated ones. The observation frequency of a topology in nature is represented by the number of homologous groups (superfamily) having the topology (See Methods for details). We regarded topologies with the observation frequency less than 1/4, at which the slope changes significantly, as unobserved. **c**, The ratio of frustration-free and frustrated β-sheet topologies depending on a degree of the observation frequency, for each number of constituent β-strands in a β-sheet. The frustration-free topologies are shown as dark gray bands and frustrated topologies, as light gray bands; the number in the bands indicates the number of each topology. The observation frequency is discretized by logarithm to base 4. **d**, The distributions of frustration-free and frustrated topologies in nature for all possible 96 topologies for 4-stranded β-sheets. Horizontal and vertical axes represent the order and orientations of the constituent β-strands, respectively. In each grid cell, a β-sheet topology is illustrated with its observation frequency in nature, which is shown by the number immediately below the topology illustration and by a gradation from white to yellow in the cell background (white is low, and yellow is high). The frustration-free and frustrated topologies are respectively colored by dark gray and light gray. The β-sheet topologies corresponding to the Greek-key and its circular permutations are marked by an asterisk. Red-color loops represent the ones including at least one frustration. Cells enclosed with a bold black square, numbered from one to eight, show unobserved frustration-free β-sheet topologies.

### Connection overlap rule

Geometrical overlap between the connections of two β-X-β motifs is disfavored (Fig. 2b). Para-β-X-β motifs have the right-handed connection preference by Richardson’s rule (Fig. 1b). Analysis of anti-β-X-β motifs in naturally occurring protein structures revealed that the connections in anti-β-X-β motifs with the jump-distance number of one have the preference of right-handed bending orientation (Extended Data Fig. 1). These right-handed connection preferences lead to the connection overlap rule (Fig. 2b).

### Connection ending rule

When the second strands in two para-β-X-β motifs are adjacent with each other and parallelly aligned, the β-sheet topologies with the S-type, in which the two connections end on the same β-sheet side, are preferred to those with the D-type, in which the two connections end on the different β-sheet sides (Fig. 2c). Analysis of para-β-X-β motifs revealed that the register shifts between the second strand in a para-β-X-β motif and the adjacent parallelly aligned β-strands are almost always zero or positive (Extended Data Fig. 2). In addition, we previously described the αβ-rule: the C_α_-C_β_ vector of the first strand residue immediately after the loop connecting from the helix to the strand points away from the helix^9^. These two preferences lead to the rule (Fig. 2c and Extended Data Fig. 3).

## Predicted novel αβ-folds with a four-stranded β-sheet

Using the set of rules for the β-sheet topology, we classified all of the open β-sheet topologies with three to eight strands into frustration-free ones without violations of the rules (these are regarded as possible topologies) and frustrated ones containing the violations. We found that many of the observed αβ-folds were identified as frustration free, while most of the unobserved αβ-folds including scarcely observed ones, as frustrated (Fig. 3a and b, see Methods). Moreover, the frustration-free β-sheet topologies were observed in more number of homologous groups (that is, evolutionary independent groups, which are referred to as superfamilies in SCOP^2^ and CATH^3^) compared with the frustrated ones (Fig. 3c, see Methods). These results suggest the capability of the set of rules to distinguish possible β-sheet topologies among all patterns of the β-sheet topologies.

We illustrated the 96 patterns of the frustrated and frustration-free β-sheet topologies for the 4-stranded αβ-proteins in Fig. 3d. Light-grey and dark-grey topologies in cells represent frustrated and frustration-free ones, respectively. The number shown immediately below the topology illustration in each cell shows the observation frequency in nature, and the cell background color also represents the observation frequency with a colour gradation from white (none) to yellow (abundant). About half of the topologies (53 patterns) are frustrated, and 37 of them are either unobserved or very rare in nature. For example, the frustrated topology located in the cell, 1342 (strand order) - ↑↓↓↓ (strand direction), violating the connection jump-distance and connection overlap rules (the violations are indicated by red color loops), is not observed in nature at all. In contrast, another half (43 patterns) are frustration-free β-sheet topologies, and 35 of them are observed in nature. For example, the frustration-free β-sheet topology located in the cell 1234 - ↑↓↑↓, called ‘meander,’ is the most frequently observed one. Here, we identified the eight frustration-free β-sheet topologies that are not observed or very rare in nature. We regarded the αβ-folds with these β-sheet topologies as possible, and attempted to carry out *de novo* design for all of the folds. Note that the β-sheet topology “8” consisting of parallelly aligned β-strands with the 3142 strand order forms a knot, which has been known as an unobserved β-sheet topology in nature and has long been considered to be impossible to exist^20,21^, but we selected this one.

## *De novo* design of all the predicted novel 4-stranded αβ-folds

To critically test whether the novel αβ-folds we predicted can be created or not, we performed the *de novo* design of αβ-fold proteins with all the predicted eight novel 4-stranded β-sheet topologies (Fig. 4a, b). Each fold was named from NF1 to NF8, according to the order of observation frequency. The folds from NF1 to NF4 are scarcely observed, and the remaining ones from NF5 to NF8 have never been observed in nature (NF6-8 have been reported as unobserved ones^21^). We sought to design these novel αβ-folds with ideal and simple structures, in which the secondary structures do not have β-bulges or α-helix kinks, and the X region in para-β-X-β motifs is an α-helix. For each novel αβ-fold, we built a backbone blueprint, in which secondary structure lengths and loop ABEGO torsion patterns are specified using the backbone design rules^9,10^ so that the target fold is favored (Fig. 4b). For NF1, 3, 4, 5, and 7, α-helices were appended at the termini to make sufficiently large hydrophobic cores. For NF5, 6, and 7, the X region in the anti-β-X-β motifs was built with an α-turn motif^22^, not just with a single loop, for the same reason; especially for NF7, ‘AAAB’ loops for βα-connections with the right twist angle (Extended Data Fig. 9) and ‘BA’ loops for αβ-connections (Extended Data Fig. 8) were adopted for making two α-turns packed together. For NF8 of a knot-forming fold, the two backbone blueprints were built using different torsion types for the loop immediately before the last strand (Extended Data Fig. 4). Next, for each blueprint, we built a backbone structure, which was obtained by averaging over several hundreds of backbone structures generated by Rosetta fragment assembly simulations^23,24^ (Fig. 4c, see Methods for details). The backbone structures were confirmed as novel ones by the database analysis using TM-align^25^ and MICAN^26,27^, and by visual inspection with the TOPS diagram^28^ (Extended Data Fig. 5). Subsequently, we carried out Rosetta design to build sidechains on each of the generated backbone structures^8,29^ (see Methods for details). Designs with low energy, tight core packing, high compatibility between local sequence and structure^9^ were selected, and their energy landscapes were explored by Rosetta ab initio structure prediction simulations^23^. Finally, designs with amino acid sequences exhibiting funnel-shaped energy landscapes toward the designed structure were characterized by experiments (Fig. 4d).

**Fig. 4.**
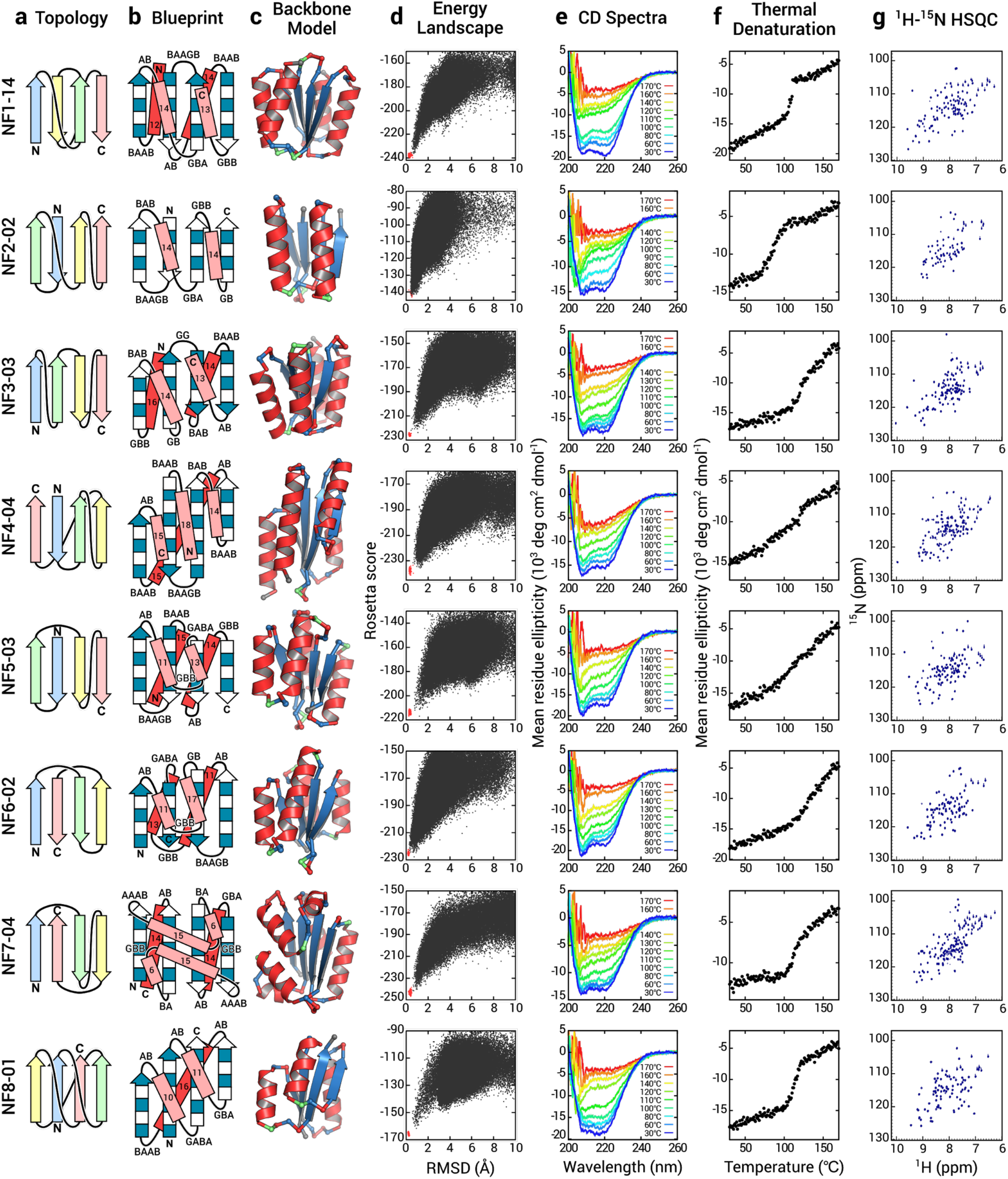
Characterization of designs for all the eight novel αβ-folds. **a**, Identified novel β-sheet topologies. **b**, Backbone blueprints used for *de novo* design of the novel αβ-fold structures. Strand lengths are represented by filled and open boxes. The filled boxes represent pleats coming out of the page, and the open boxes represent pleats going into the page. The letter strings next to the loops indicate their ABEGO torsion patterns^10^. **c**, Backbone structures generated from the blueprints. Each residue color represents its ABEGO torsion angle (red: A, blue: B, and green: G). **d**, Energy landscapes obtained from Rosetta *ab initio* structure prediction simulations. Each point represents the lowest energy structure obtained in an independent trajectory starting from an extended chain (black) or the design model (red) for each sequence; the x-axis shows the Cα root mean square deviation (RMSD) from the design model and the y-axis shows the Rosetta all-atom energy. **e**, Far-ultraviolet circular dichroism (CD) spectra at various temperatures (30-170 °C). **f**, Thermal denaturation monitored at 222 nm. **g**, Two-dimensional ^1^H-^15^N HSQC spectra at 25 °C and 600 MHz.

## Experimental characterization

We obtained synthetic genes encoding 16 designs for NF1, 4 for NF2-3, 6 for NF4-7, and 12 for NF8 (6 for each of the two different blueprints) (all these sequences are described in Supplementary Tables 1-8). For all of the sequences, no clear homologous proteins to any known protein were found (All designs have BLAST *E*-value >10 ^-3^ against the NCBI nr database of non-redundant protein sequences). The proteins were expressed in *E. Coli* and purified by a Ni-NTA affinity column. For all target folds, 56 of 60 designed proteins were found to be expressed well and soluble. These were then characterized by circular dichroism (CD) spectroscopy, size-exclusion chromatography combined with multi-angle light scattering (SEC-MALS), and ^1^H-^15^N heteronuclear single quantum coherence (HSQC) NMR spectroscopy. The experimental results for all designs for all target folds are summarized in Extended Data Table 1. The success rate of the designs including the knotted fold was as high as those in the previous *de novo* designs with the folds widely existing in nature (28 of 60 designs were characterized as foldable proteins)^9–13,30^. For each target fold, one monomeric design with the CD spectrum characteristic of αβ-proteins and the expected number of well-dispersed sharp NMR peaks were selected for NMR structure determination (Fig. 4e-g). The NMR structures solved by using MagRO-NMRViewJ^31,32^ were in close agreement with the computational design models, with the correct β-sheet topologies (Fig. 5, the root mean square deviation (RMSD) values for backbone heavy atoms were ranged from 1.4 to 2.0 Å; Extended Data Table 2 for NMR constraints and structure statistics). These results demonstrated that all the novel αβ-folds predicted by our rules can indeed be created. Remarkably, we succeeded in designing the smallest knotted NF8 structure consisting of only 4 strands (Extended Data Fig. 6).

**Fig. 5.**
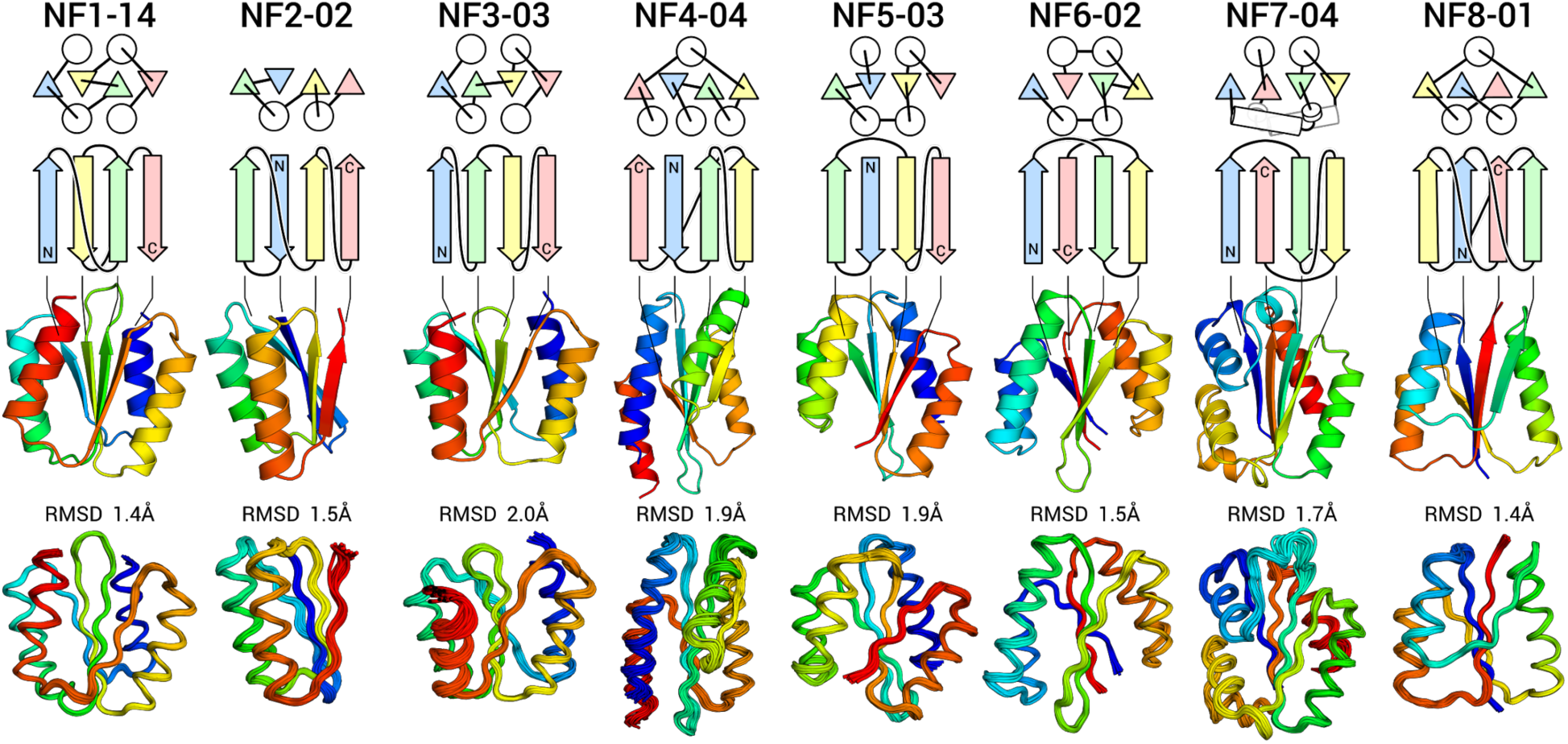
Comparison of computational models with experimentally determined structures. (Top) The designed novel αβ-folds from NF1 to NF8. The tertiary arrangement of α-helices (circles) and β-strands (triangles) and their connections are shown in the upper; the β-sheet topology is shown in the lower. (Middle) Computational design models. (Bottom) The NMR structures. The RMSD for backbone heavy atoms between the design model and NMR structures is indicated. The design models are available in Supplementary Dataset, and the NMR structures are available in PDB; NF1-14 (PDB ID, 7BPL), NF2-02 (7BPM), NF3-03 (7BQE), NF4-04 (7BQC), NF5-03 (7BPP), NF6-02 (7BQB), NF7-04 (7BPN), and NF8-01 (7BQD).

## Prediction of novel αβ-folds with five- to eight-stranded β-sheets

The success in *de novo* design of all the predicted eight novel αβ-folds demonstrated the ability of the set of rules to predict novel αβ-folds. We then revisit the number of frustration-free unobserved β-sheet topologies with five- to eight-stranded β-sheets, shown in Fig. 3a (for 3-stranded αβ-proteins, all ten frustration-free β-sheet topologies have been observed in nature). As the number of constituent β-strands in a β-sheet increases, the number of frustration-free unobserved topologies increases exponentially and the ratio of unobserved topologies in the frustration-free ones also increases. The prediction indicates that an enormous number of frustration-free (i.e., possible) αβ-folds have been left as unobserved in nature; the number (∼12000) is far more than that of the αβ-folds observed in nature (400). Note that since we only investigated novel folds that are identified by our introduced set of rules, the predicted number corresponds to a lower limit of that of novel folds. There must be more novel folds that are not identified by the rules but accessible to polypeptide chains.

## Discussion

How large the protein fold space is that is accessible to the polypide chain has been unknown. We systematically investigated the unexplored fold space by introducing a set of rules to predict novel αβ-folds, and by carrying out *de novo* design of all the predicted novel αβ-folds with a four-stranded β-sheet. As the results, we found that all the predicted novel αβ-folds, including a knotted fold, can be created. Remarkably, the design success rate was comparable to that of previous *de novo* designs with naturally occurring folds, and the thermal stability of the designs was as high as previous designs^9–12,30,33^. Our study indicates that there are at least about twelve thousand novel αβ-folds with five- to eight-stranded β-sheets.

The possible reasons that the novel folds have not been reached by natural evolution can be considered in the following. 1) The folds in nature have been repetitively reused and adapted for different functions, and the novel folds have not been employed just by accident, 2) the time scale of biological evolution so far can be too short to sample all novel folds, and 3) the novel folds are incapable of carrying out functions required by life. To address such questions, the relations between novel fold structures and their functions need to be studied.

The number of predicted novel αβ-folds, which is at the lower limit of that of novel αβ-folds, is far more than that of the observed folds in nature. Moreover, the novel αβ-folds include knot-forming ones. Recently, functional proteins have been designed *de novo*^14,34–44^. Our predicted novel αβ-folds provide an enormous scaffold set for designing protein structures with desired functions. We are at a great starting point for exploring the universe of protein structures beyond natural evolution.

## Methods

### Structure dataset of naturally occurring proteins

For the derivation of a set of rules for the β-sheet topology (Fig. 2), a dataset containing 12,595 chains obtained from the cullpdb database (date; 12/13/2018)^45^ with more than 40 residues, sequence identity < 25%, resolution < 2.5 Å, and R-factor < 1.0 was used. For the analysis of β-sheet topologies of naturally occurring protein structures (Fig. 3), a dataset containing 65,371 domains obtained from a semi-manually curated domain database ECOD^46^ (This database provides a hierarchical grouping of evolutionarily related domains) with more than 40 residues and sequence identity <99% was used. Structure refinements were carried out by ModRefiner^47^ for all obtained structures, and then the secondary structures were assigned by STRIDE^48^; when the Cα RMSD of a refined structure against the original structure was greater than 1.0 Å, the refined structure was discarded and the original one was used.

### Analysis of β-sheet topologies in naturally occurring proteins

β-sheet topologies were defined for open β-sheets included in the protein domains obtained from ECOD, with the following criteria: 1) the lengths of constituent β-strands are more than two residues, 2) the number of β-strands is at least three, 3) two neighboring β-strands have at least two main-chain hydrogen bonds between the β-strands, and 4) no insertion along a sequence by any β-strands belonging to another β-sheet consisting of more than two β-strands (Extended Data Fig. 7). Additionally, “branched’’ β-sheets that include β-strands having more than two neighboring β-strands were discarded.

Frequencies of all β-sheet topologies in nature were studied using the ECOD database, in which protein domains are classified according to their evolutionary relationships. In the database, the two categories, Family and Homology, are defined. Family represents a group of evolutionarily related protein domains, identified by substantial sequence similarity, and Homology represents a group comprising multiple Family groups, of which evolutionary relationships are inferred on the basis of functional and structural similarities (Homology is equivalent to the superfamily in the SCOP^2^ or CATH^3^ structure databases). To study the observation frequency for each β-sheet topology, we counted the number of Homology groups having the topology, with the following consideration. We first examined the occupation ratio of the topology in the *i*-th Homology group:

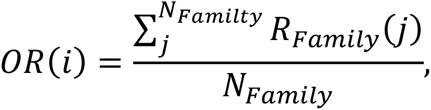

where *N*_Family_ is the total number of Family groups belonging to the Homology group, and *R*_Family_*(j)* is the ratio of protein domains having the β-sheet topology in the *j*-th Family group. Thus, when all domains in the Homology group contain the β-sheet topology, the occupation ratio of the Homology group is one, otherwise less than one. Finally, the observation frequency for each topology is calculated by the sum of the occupation ratios across Homology groups,

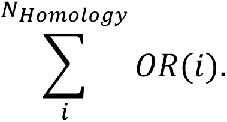

For four-stranded β-sheet proteins, we manually checked all structures having the topologies of which the observation frequencies are less than 1.0, and then changed the β-sheet assignments for some of the structures: the β-sheets included in e3hy2X1, e1xw3A1, e2hwjA2, e4rsfA1, and e1tocR2 were identified as β-barrel, and those of e4rgzA1, e2bjjX3, e1iejA2, e3s9lC3, e1blfA4, and e2d3iA3 were, as 6-stranded β-sheet.

### Backbone construction

We built a backbone blueprint for each novel αβ fold. For the X region in para-β-X-β motifs, a helix was built. The lengths of secondary structures and the ABEGO torsion patterns for the connecting loops were obtained from the design rules described in previous papers^10^. For NF1, 3, 4, 5, and 7, α-helices were appended to their termini to make a sufficiently large hydrophobic core between the β-sheet and the α-helices. For the same purpose, for the X region in antiparallel β-X-β motifs in NF5, 6, and 7, an α-turn structure consisting of a helix-loop-helix unit was built. β-strand lengths were chosen from four to seven, and α-helix lengths were from eleven to seventeen residues. The torsion ABEGO patterns of loops were determined by referring to the previous paper^10^: GB, GBA, or BAAB for the connecting loops of the para type αβ-unis, AB for the para type βα-units, BAB,or GBB for the anti type βα-units, BAAGB for the R-chirality ββ-units and GG for the L-chirality ββ-units. In this paper, we newly introduced GABA for the para type αβ-units and GBB for anti type αβ-units (Extended Data Fig. 8). For the α-turn structures, we used the GBB loop^22^ for connecting the two helices. In the NF7 blueprint, AAAB for the anti type βα-units (Extended Data Fig. 9) and BA for the para type αβ-units (Extended Data Fig. 8) were used to arrange the two α-turns packed with each other.

1,000-40,000 backbone structures for each blueprint (sufficient number depends on its fold type) were generated were generated by Rosetta sequence-independent Monte-Carlo fragment assembly simulations using a coarse-grained model backbone structures of backbone structures, in which each residue is represented by mainchain atoms (N, H, CA, C and O) and a side-chain pseudo atom ^49,50^. The Rosetta potential function used in the simulations includes steric repulsion (vdw = 1.0), overall compaction (rg = 1.0), secondary structure pairings (ss_pair = 1.0, rsigma = 1.0, and hs_pair = 1.0), and main-chain hydrogen bonds (hbond_sr_bb = 1.0 and hbond_lr_bb = 1.0), with no sequence dependent score terms. The steric radius of Val was used for that of the side-chain pseudo atom. The ss_pair and rsigma score terms are modified so that only the strand residue pairs, specified in the blueprint, are favored in the simulations. To enhance the sampling efficiency for obtaining target topology backbone structures, we built backbone structures parts by parts. For instance, in the case of the NF2 fold, the N-terminal half (β1-β2-α1-β3), which forms a locally globular substructure, was built first, and successively, the C-terminal half (α2-β4) was built by extending the N-terminal half. The generated backbone structures were further refined by the following two steps. 1) β-sheet refinement. The entire structures were minimized with the constraints making the Cα atoms of the neighboring strand-residues in the blueprint being less than 5.5 Å, using the Rosetta full-atom FastRelax protocol with the upweighted hydrogen-bonding and a backbone torsion angle terms (hbond_sr_bb = 5.0, hbond_lr_bb = 3.0, and omega = 3.0). Val was used for the full-atom side-chains for all residues except for those at the G region in the ABEGO Ramachandran map (Gly was used). This step was repeated up to 10 times until the secondary structures and ABEGO torsion patterns became identical to those designated in the blueprint. (2) α-helix refinement. The loop-helix-loop structures were rebuilt using the CCD loop closure method implemented in the BlueprintBDR mover. This step was repeated up to 10 times for each loop-helix-loop region until the α-helix was built without kinks and the loop torsion patterns were identical to those designated in the blueprint. Next, we selected 100-500 backbone structures, in which the terminal α-helices are packed with the β-sheet; we required that at least one residue in any continuous five residue segments in the terminal α-helices to be buried (the accessible surface area < 40 Å^2^) with the central two β-strands in the β-sheet. The generated backbone structures sometimes show structural diversity. In such cases, we clustered the backbone structures based on structure similarity, using a hierarchical clustering approach (average linkage). The structural similarity was evaluated with Cα RMSD, and the cutoff for the clustering was determined within a range from 1.0 to 2.0 Å, according to the structural diversity of the generated structures. From the top three largest clusters, we selected the cluster consisting of the structures with tightly packed secondary structures. Finally, we averaged xyz-coordinates of the main-chain atoms of 30-150 backbone structures in the cluster, followed by the Rosetta Idealization protocol with the upweighted score terms (hbond_sr_bb = 10.0, hbond_lr_bb = 10.0, and omega = 10.0), resulting in a backbone structure to be used for the following side-chain design.

### Sequence design

We performed RosettaDesign calculations^51^ with the full-atom Talaris2014^49^ scoring function to design side-chains (amino acid sequences) that stabilize each generated backbone structure. The design calculation consists of the following three steps. 1) Several cycles of amino-acid sequence optimization with a fixed backbone and the following backbone relaxation. 2) Mutations of buried polar residues to hydrophobic ones, followed by the entire structure optimization. 3) Mutations of solvent-exposed hydrophobic residues to polar residues, followed by the entire structure optimization. Amino-acid residue types to be used for the design of each residue position except for that of loop regions were restricted based on the secondary structure of the position and the buriedness calculated using virtual amino acids. For the design of each loop region (the residues in the loop and the preceding and following three residues), amino acid types were restricted on the basis of the consensus amino acids obtained from the sequence profile for naturally occurring protein structure fragments, which were collected using the following criteria, 1) identical secondary structure and ABEGO torsion pattern to the loop region, and 2) RMSD against the loop structure is lower than 2.0 Å. Through the RosettaDesign calculations, up to 40,000 designs were generated for each design target structure.

The designed sequences were then filtered based on the Rosettatotal energy, the RosettaHoles score^52^ < 2.0, and the packstat score^52^ > 0.55 for the target NF2 and > 0.6 for others. Furthermore, we filtered the designs on the basis of the local sequence-structure compatibility^53^. We collected 200 fragments for each nine-residue frame in each designed sequence from a non-redundant set of experimental structures, based on the sequence similarity and secondary structure prediction. Subsequently, for each frame, we calculated Cα RMSD of the local structure against each of the 200 fragments. Designs were ranked according to the summation of the log-ratio of the fragments, for which the RMSD was less than 1.5Å, across all nine-residue frames, and those with high values were selected.

### Protein expression and purification

A spacer was added at the C-terminus of each designed sequence (’GSWS’ for the sequences that have neither a TRP residue nor more than two TYR residues and ’GS’ for others) to separate the designed region from the C-terminal 6xHis-tag. The genes encoding the designed sequences, which were cloned into pET21b expression vectors, were synthesized by Eurofins Genomics (Tokyo, Japan). The designed proteins were expressed in *E. coli* BL21 Star (DE3) cells (Invitrogen) as uniformly (U-)^15^N-labeled proteins using MJ9 minimal media^54^, which contain ^15^N ammonium sulfate as a sole nitrogen source and ^12^C glucose as a sole carbon source, respectively. The expressed proteins with a C-terminal 6xHis-tag were purified through a Ni-NTA affinity column. The purified proteins were then dialyzed against typical PBS buffer, 137 mM NaCl, 2.7 mM KCl, 10 mM Na_2_HPO_4_, and 1.8 mM KH_2_PO_4_, at pH 7.4; this buffer was used for all the experiments except NMR structure determination. The expression, solubility, and purity of the designed proteins were assessed by SDS-PAGE and mass spectrometry (Thermo Scientific Orbitrap Elite). The protein concentrations were determined from the absorbance at 280 nm^55^ using an ultraviolet spectrophotometer (NanoDrop, Thermo Scientific).

### Circular dichroism (CD) spectroscopy

All CD data were collected on a JASCO J-1500 KS CD spectrometer. For all designs, far-UV CD spectra were measured from 260 to 200 nm using ∼20 μM protein samples in PBS buffer (pH 7.4) with a 1 mm path length cuvette. For the representative eight designs (NF1-14, NF2-02, NF3-03, NF4-04, NF5-03, NF6-02, NF7-04, and NF8-01), thermal denaturation measurements were performed from 30 to 170 °C under 1 MPa pressure with an increase of 1 °C per minute. During the denaturation, the ellipticity at 222 nm was monitored, and far-UV CD spectra were measured from 260 to 200 nm at various temperatures shown in Fig. 4e.

### Size-exclusion chromatography combined with multi-angle light scattering (SEC-MALS)

SEC-MALS experiments were performed by using a miniDAWN TREOS static light scattering detector (Wyatt Technology Corp.) combined with an HPLC system (1260 Infinity LC, Agilent Technologies). The volume 100 μl of 200-500 μM protein samples in PBS buffer (pH 7.4) after Ni purification was injected into a Superdex 75 Increase 10/300 GL column (GE Healthcare) or a Shodex KW-802.5 (Showa Denko K.K.) equilibrated with PBS buffer at a flow rate of 0.5 mL/min. The protein concentrations were calculated from the absorbance at 280 nm detected by the HPLC system. Static light scattering data were collected at three different angles 43.6°, 90.0°, and 136.4° at 659 nm. These data were analyzed with ASTRA software (version 6.1.2, Wyatt Technology Corp.) with a change in the refractive index with concentration, a *dn/dc* value, 0.185 ml/g.

### 2D ^1^H-^15^N HSQC measurement by nuclear magnetic resonance (NMR)

2D ^1^H-^15^N HSQC spectra were measured to verify whether the designed proteins fold into well-packed structures. The HSQC spectra were collected for 0.5-1.0 mM protein samples in 90% ^1^H_2_O/10% ^2^H_2_O PBS buffer (pH 7.4) at 25 °C on a JEOL JNM-ECA 600 MHz spectrometer. The stable monomeric design with a most well-dispersed and sharp NMR spectrum for each fold (NF1-14, NF2-02, NF3-03, NF4-04, NF5-03, NF6-02, NF7-04, and NF8-01) was selected for NMR structure determination.

### Determination of solution structure by NMR

#### Sample preparation

For the NMR structure determination of the eight selected designs, the uniformly isotope-labeled [U-^15^N, U-^13^C] proteins were expressed using the same method as described above except ^13^C glucose as a sole carbon source. The U-^15^N, U-^13^C-enriched proteins were purified through Ni-NTA affinity column and further by gel filtration chromatography on an ÄKTA Pure 25 FPLC (GE Healthcare) using Superdex 75 Increase 10/300 GL column (GE Healthcare). The purified proteins were dissolved in 95% ^1^H_2_O/5% ^2^H_2_O PBS buffer at various pH; (50 mM NaCl, 1.1 mM Na_2_HPO_4_, and 7.4 mM KH_2_PO_4_ at pH 6.0) for NF2-02, NF3-03, and NF6-02, (50 mM NaCl, 4.3 mM Na_2_HPO_4_, and 5.7 mM KH_2_PO_4_ at pH 6.8) for NF5-03, NF7-04, and NF8-01, (50 mM NaCl, 5.6 mM Na_2_HPO_4_, and 1.1 mM KH_2_PO_4_ at pH 7.4) for NF1-14, (137 mM NaCl, 1.1 mM Na_2_HPO_4_, and 7.4 mM KH_2_PO_4_ at pH 7.4) for NF4-04, respectively. Shigemi micro NMR tubes were used for all NMR measurements except residual dipolar coupling (RDC) (protein concentration ∼900 μM for all designed proteins except NF4-04 and NF6-02; ∼400 μM for NF4-04; ∼700 μM for NF6-02), and normal NMR tubes were used for RDC experiments (protein concentration ∼200 μM).

#### NMR measurements

NMR measurements were performed on Bruker AVANCE III NMR spectrometers equipped with QCI cryo-Probe (^1^H/^13^C/^15^N/^31^P) at 303 K. The spectrometers with 600, 700, and 800 MHz magnets were used for the signal assignments and NOE related measurements, while 900 and 950 MHz ones, for RDC experiments. For the signal assignments, 2D ^1^H-^15^N HSQC (echo/anti-echo), ^1^H-^13^C Constant-Time HSQC for aliphatic and aromatic signals, and 3D HNCO, HN(CO)CACB, and 3D HNCACB for backbone signal assignments, were measured, while BEST pulse sequence^56^ was applied to the triple resonance measurements for NF2, NF3, NF5, NF6, NF7, and NF8. For structure determination, 3D ^15^N-edited NOESY, 3D ^13^C-edited NOESY for aliphatic and aromatic signals (mixing time = 100 ms) were performed. For RDC experiments, 2D IPAP ^1^H-^15^N HSQC spectra using water-gate pulses for water suppression were measured with or without 6-10 mg/ml of Pf1 phage (ASLA biotec Ltd.). For confirming the positions of ^1^H-^15^N signals in the 2D IPAP ^1^H-^15^N HSQC, 3D HNCO at the identical buffer condition containing Pf1 phage were measured. The α- and β-states of ^15^N signals split by ^1^H-^15^N ^1^J-coupling were separately identified for the protein in the isotropic and weakly aligned states, in order to obtain 1-bond residual dipolar coupling 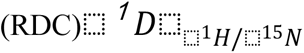 values. They were estimated by simple subtraction of the shifted values between isotropic and weakly aligned states then divided by the static magnetic field to get the RDC value in Hz.

#### NMR signal assignments

All NMR signals were identified by using MagRO-NMRViewJ (upgraded version of Kujira^31^) in a fully automated manner, then noise peaks were filtered by deep-learning methods using Fit_Robot^32^. FLYA module was used for fully automated signal assignments and structure calculation^57^ to obtain roughly assigned chemical shifts (Acs) then the trustful ones were selected into MagRO Acs table. After confirmation and correction of the Acs by visual inspection on MagRO, TALOS+^58^ calculations were performed to predict phi/psi dihedral angles, which were then converted to angle constraints for CYANA format. The signal assignments in 2D ^1^H-^15^N HSQC spectrum for each fold are shown in Supplementary Figs. 1-8.

#### Structure calculation

Several CYANA^59^ calculations were performed using the Acs table, NOE peak table, and dihedral angle constraints. After the CYANA calculations, several dihedral angle constraints derived from TALOS+ revealing large violations for nearly all models in the structure ensemble were eliminated. After the averaged target function of the ensemble reached less than 2.0 Å^2^, refinement calculations by Amber12 were carried out for 20 models with the lowest target functions.

#### NMR structure validation

The RMSD values were calculated for the 20 structures overlaid to the mean coordinates for the ordered regions, automatically identified by Fit_Robot using multi-dimensional non-linear scaling^60^. The RDC back-calculation was performed by PALES^61^ using experimentally determined values of RDC. The averaged correlation between the simulated and experimental values was obtained using the signals except for the residues on overlapped regions in ^1^H-^15^N HSQC, and for the residues predicted to be low order parameters (S^2^ < 0.8) by TALOS+. The detailed methods and results are described in Extended Data Table 2 and Supplementary Document.

## Acknowledgments

We thank RIKEN Yokohama NMR Facility for NMR spectra measurements, Functional Genomics Facility, NIBB Core Research Facilities, especially Y. Makino, for mass spectrometry analysis, and Instrument Center for HSQC spectra measurements, Okazaki, Japan. We also thank M. Yamamoto, Naoya Kobayashi, and M. Kondo for support and discussion of experiments; K. Sakuma for discussion on computational design; T. Kosugi for support and discussion of experiments, and constructive discussion on the manuscript; M. Sasai, M. Ota, and D. Baker for valuable comments and discussion on the manuscript. The computations were performed using the Research Center for Computational Science (RCCS), Okazaki, Japan. NMR structure determination was supported by Basis for Supporting Innovative Drug Discovery and Life Science Research (BINDS) from AMED under Grant Number JP21am0101072. This work was supported by the Astrobiology Center Program of National Institutes of Natural Sciences (NINS) (Grant Number AB291007) and the Japan Society for the Promotion of Science (JSPS) KAKENHI Grants-in-Aid for Scientific Research 15H05592 to N. Koga, 18H05420 to N. Koga, and 19H03166 to G. Chikenji and N. Koga, BINDS from AMED under Grant Number JP20am0101111 to G. Chikenji, and JSPS Research Fellowship (PD) 17J02339 to S. Minami.

## Author contributions

S.M., N.Kobayashi, R.K., G.C., and N.Koga designed the research; S.M. performed database analysis, design of all proteins, and experimental characterizations except for the NMR structure determination; G.C. provided a core idea for the database analysis; N.Kobayashi, T.S., and N.T. performed the NMR structure determination experiments; N.Kobayashi performed the NMR structural analysis; and S.M., N.Kobayashi., T.F., R.K., and N.Koga wrote the manuscript.

## Additional Information

**Supplementary Information** is available for this paper.

## Extended Data Figures

**Extended Data Fig. 1.**
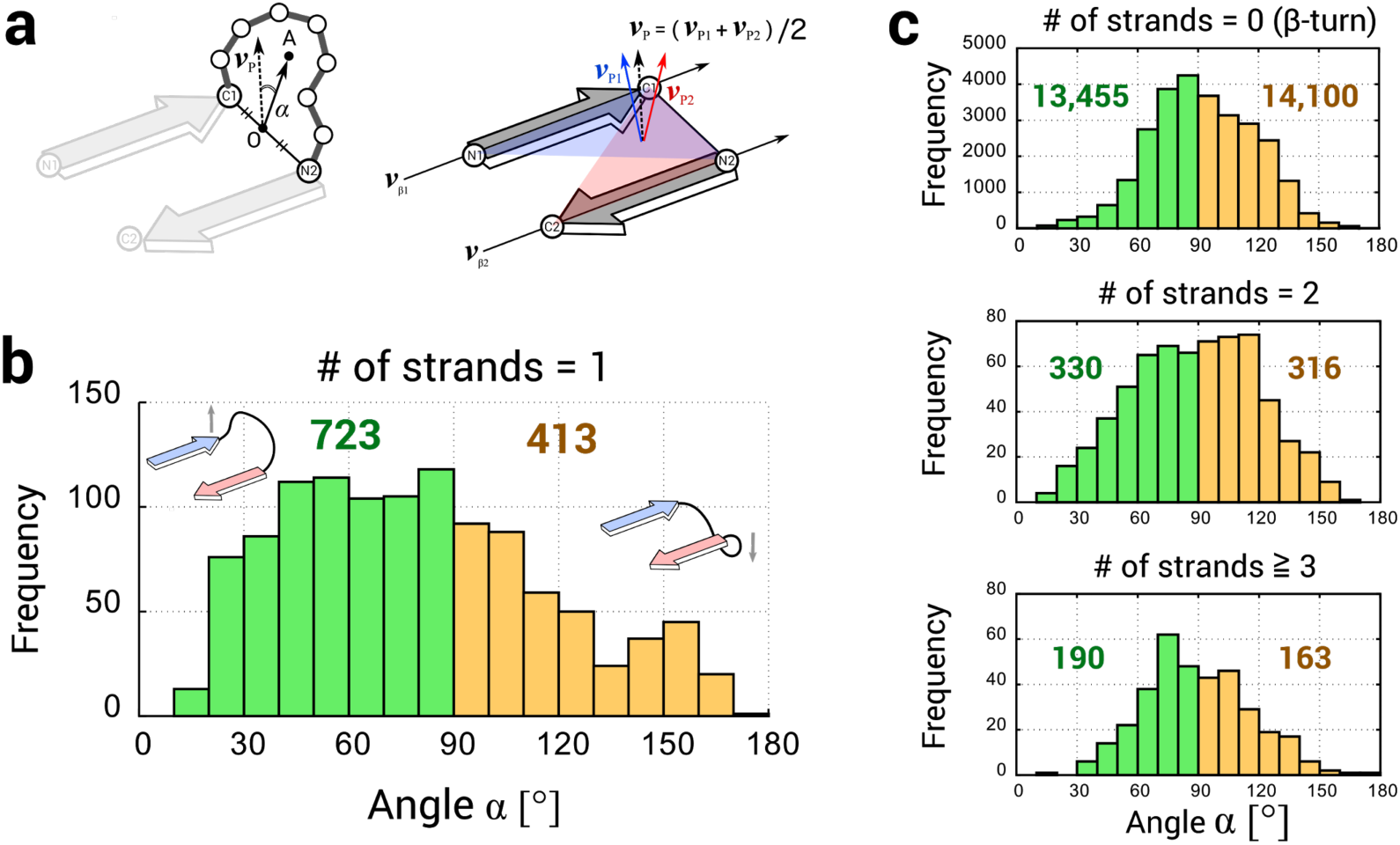
The bending orientation preference of anti-β-X-β motifs with the connection jump-distance of one. **a**, Left: the bending angle, α, for anti-β_1_-X-β_2_ motifs was identified as the angle between the β-sheet normal vector ***v***_p_ and the vector from the midpoint O of the terminal β-strand backbone atoms, C1 (carbonyl carbon of the first strand) and N2 (amide nitrogen of the second strand), to the average coordinate A over the loop Cα atoms. Right: the ***v***_p_ was calculated by averaging the normal vectors to the two planes, which are respectively defined by the following backbone atoms, N1-C1-N2 and C1-N2-C2. **b**, The distribution of the angle α for naturally occurring protein structures with the jump-distance number of one. Anti-β-X-β motifs with the bending angle less than 90° are more frequently observed than those with more than 90°, indicating the right-handed bending orientation preference of anti-β1-X-β2 motifs with the jump-distance number of one. This preference may arise from the intrinsic chirality and geometrical preferences of the polypeptide chain. **c**, The distributions of the angle α for naturally occurring protein structures with the jump-distance number of zero (top), two (middle), and larger than two (bottom), respectively. No preferences of the bending angle were observed.

**Extended Data Fig. 2.**
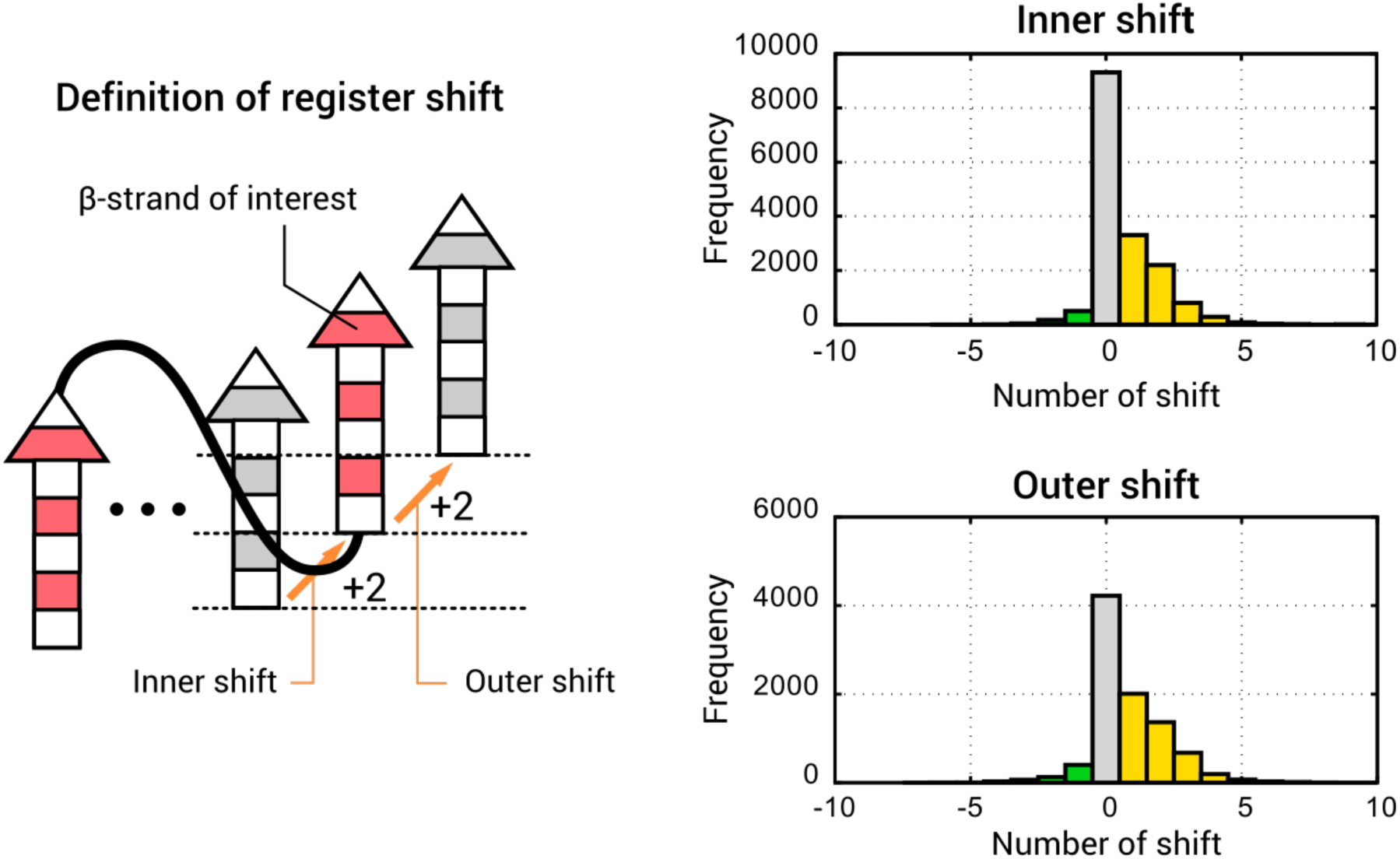
The register-shift rule for para-β-X-β motifs. The register shifts for a para-β-X-β motif are identified in the relations of the second strand (red) in the β-X-β motif with the adjacent parallelly aligned β-strands (grey): the inner shift is defined with the grey β-strand inside the para-β-X-β motif, and the outer register shift is defined with the grey β-strand outside the para-β-X-β motif. Analysis of the residue offset for the inner and outer shifts for para-β-X-β motifs in naturally occurring protein structures revealed that the register shifts are mostly zero or positive; the origin of this preference is partly explained by energetic penalties of steric repulsion and buried polar atoms that emerge when unfavored register shifts are formed.

**Extended Data Fig. 3.**
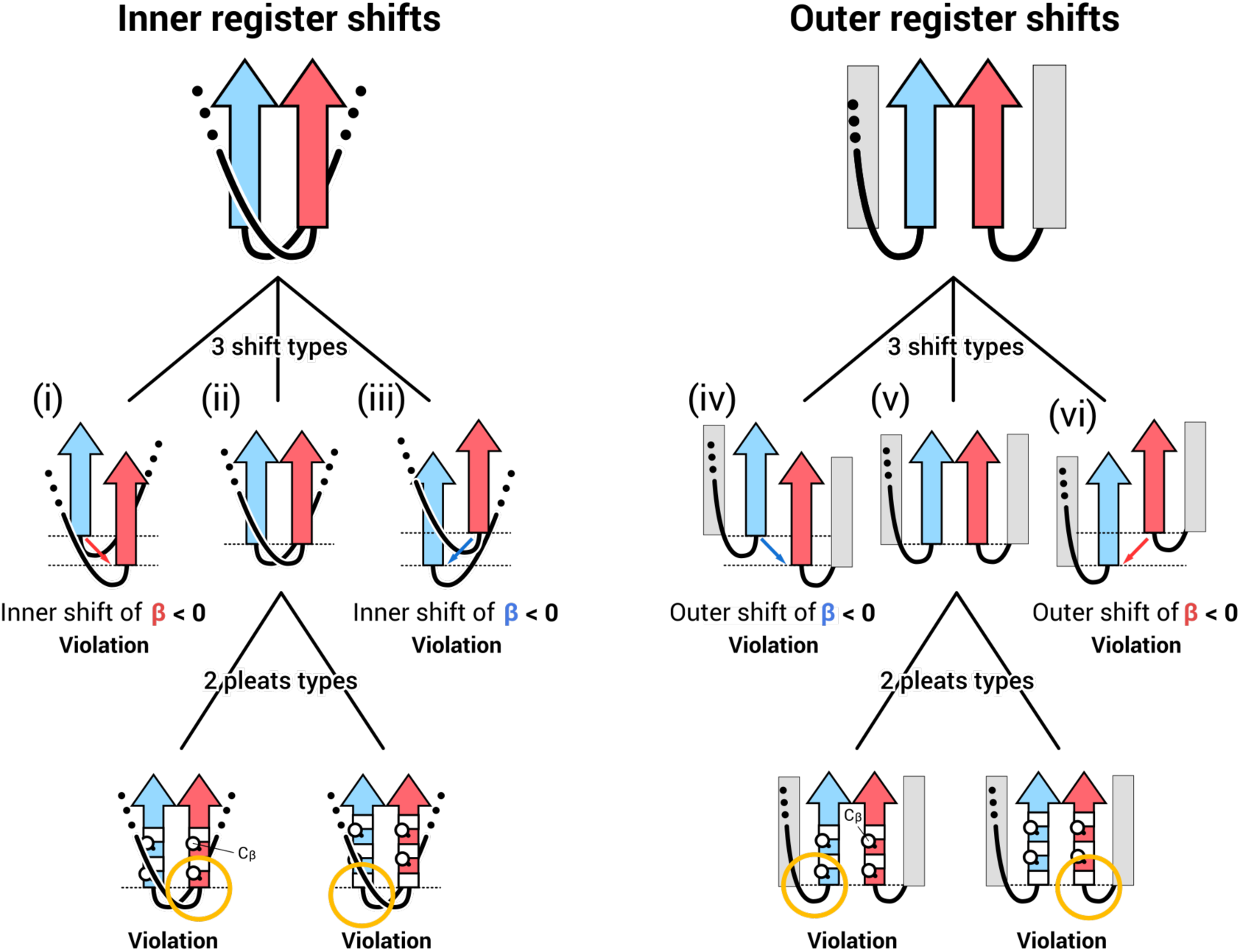
The origin of the connection ending rule. The inner and outer register shift arrangements for para-para-β-X-β motifs (red and blue), violating the connection ending rule, are illustrated in the left and right, respectively (only the second strands of the para-β-X-β motifs are shown). In the arrangements, the second strands are adjacent to each other and the connections are on the different β-sheet sides, which do not agree with either the register-shift rule (Extended Data Fig. 2) or αβ-rule^9^. In the cases of the register shift of non-zero, (i), (iii) (iv), and (vi), the β-X-β motifs violate the register shift rule (Extended Data Fig. 2). In (i) and (vi), the red strand shifted toward the negative orientation against the blue strand; in (iii) and (iv), the blue strand shifted toward the negative orientation against the red strand. In the cases of the register shift of zero, (ii) and (v), the αβ-rule^9^ is violated: the vector from the Cα to Cβ atoms of the first strand residue in either the β-strands points toward the X region in β-X-β motifs.

**Extended Data Fig. 4.**
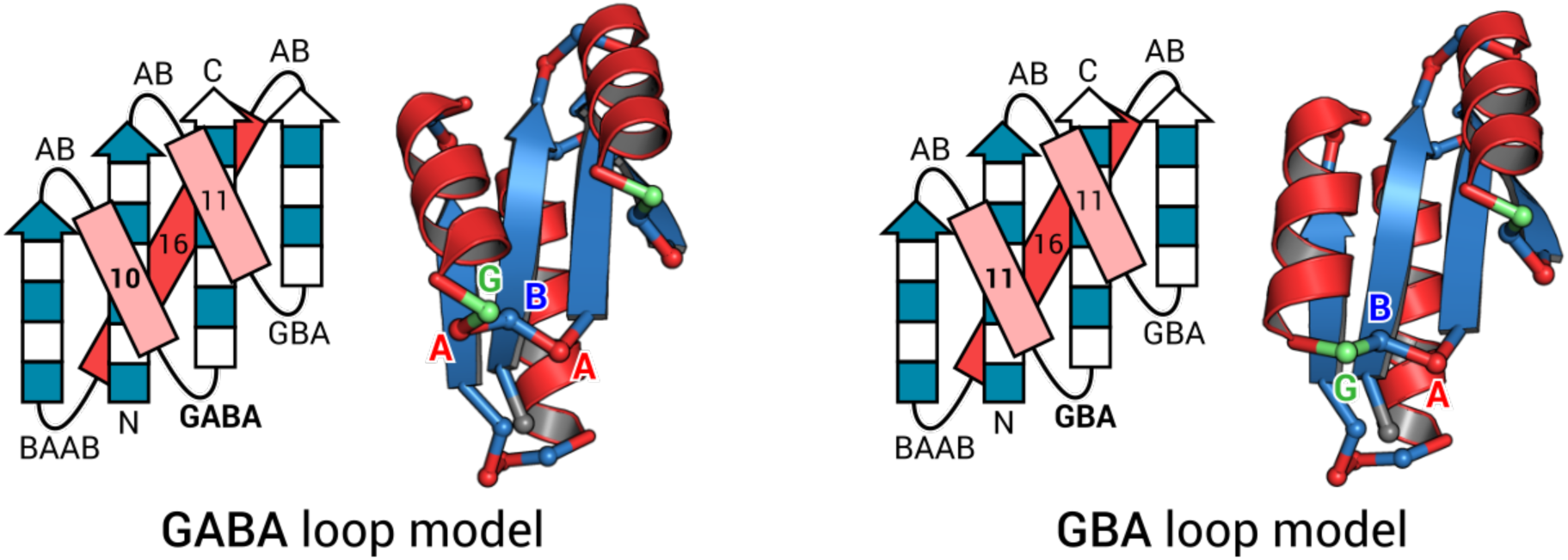
Two different backbone blueprints used for the design of the target NF8. The torsion patterns immediately before the last strand are different.

**Extended Data Fig. 5.**
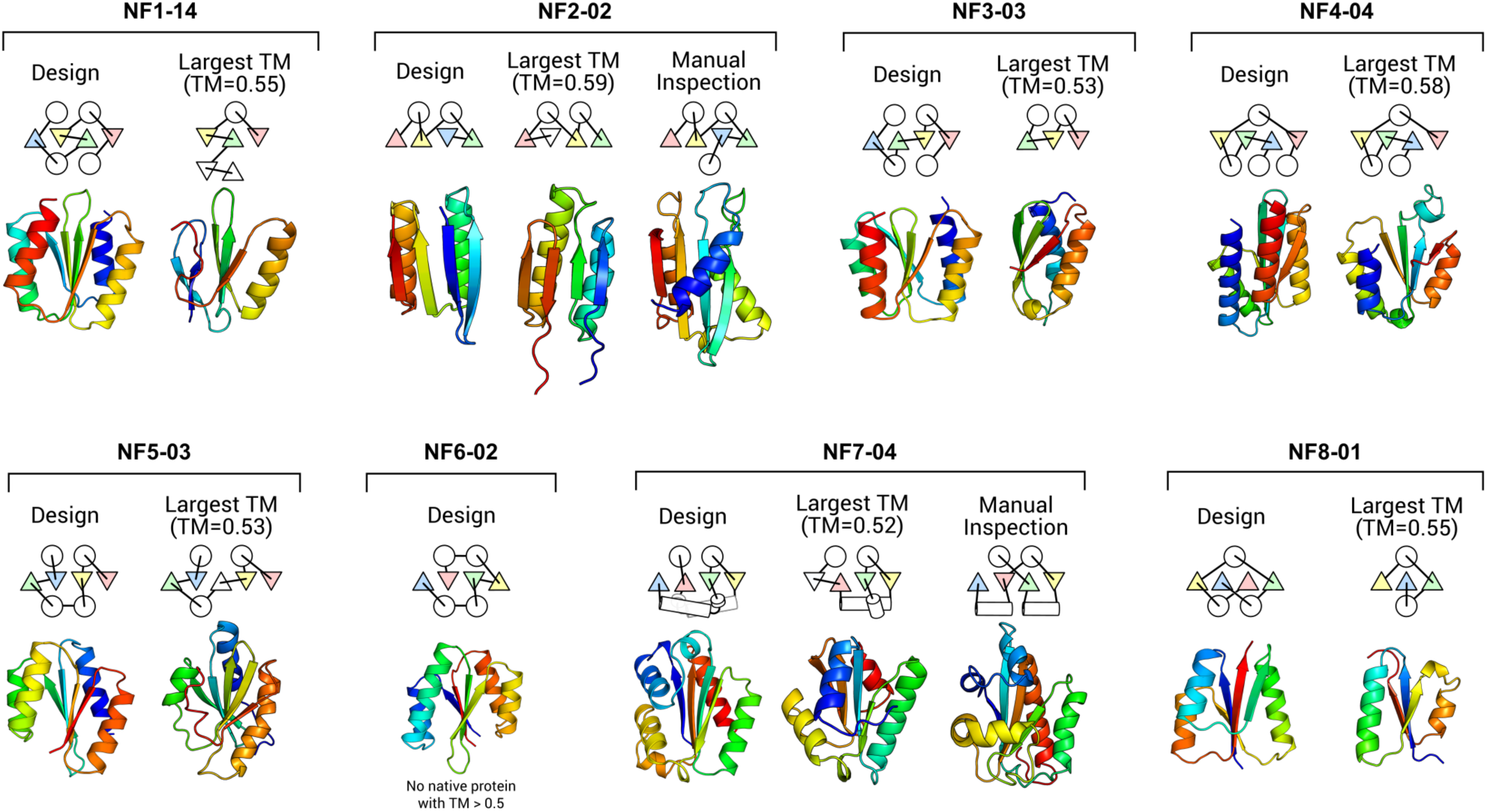
The most similar naturally occurring protein structure to each designed novel fold structure. For each designed novel fold structure, similar domain structures were searched by MICAN^26,27^ (sequential mode) and TM-align^25^ against the ECOD domain dataset (99% sequence non-redundant set). We first collected all domains with TM-score larger than 0.5 compared to each target and then examined them manually. The domain with the largest TM-score for each target is shown in each panel, except for the NF6-02 (there is no domain with TM-score larger than 0.5). No similar structures were found except for the NF2 and NF4 designs. Although these β-strand arrangements are identical to those of the corresponding design targets, these folds can be regarded as different in terms of the original fold definition: the structure similar to the NF2 design includes an additional N-terminal helix, and the one for the NF4 design does not have the C-terminal helix.

**Extended Data Fig. 6.**
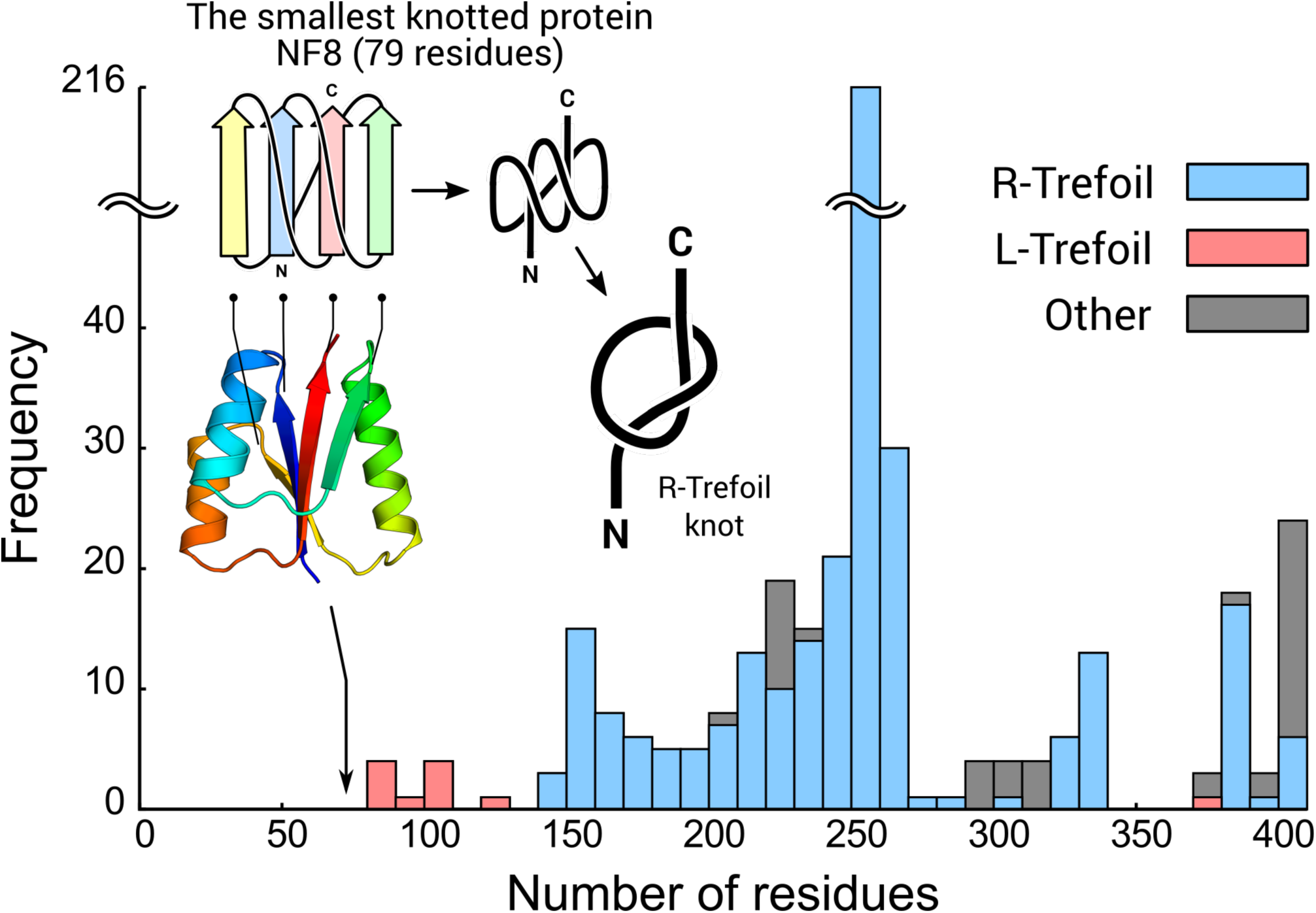
The smallest knotted protein, the design NF8. The stacked histogram represents the number of naturally occurring knot proteins in the PDB, depending on the chain length (the original annotation data was obtained from the KnotProt database^63^). The blue, red, and gray bars represent the right-handed trefoil knot (R-Trefoil), left-handed trefoil knot (L-Trefoil), and other knot types (Other), respectively. The design NF8 with the R-Trefoil knot, indicated by an arrow, is characterized as the smallest knotted protein with 79 residues. Note that this is an exceptional case for the R-Trefoil knot structures; the minimal size observed in nature is about 140 residues (For the L-Trefoil structure, the smallest one has 82 residues, PDB ID; 2EFV).

**Extended Data Fig. 7.**
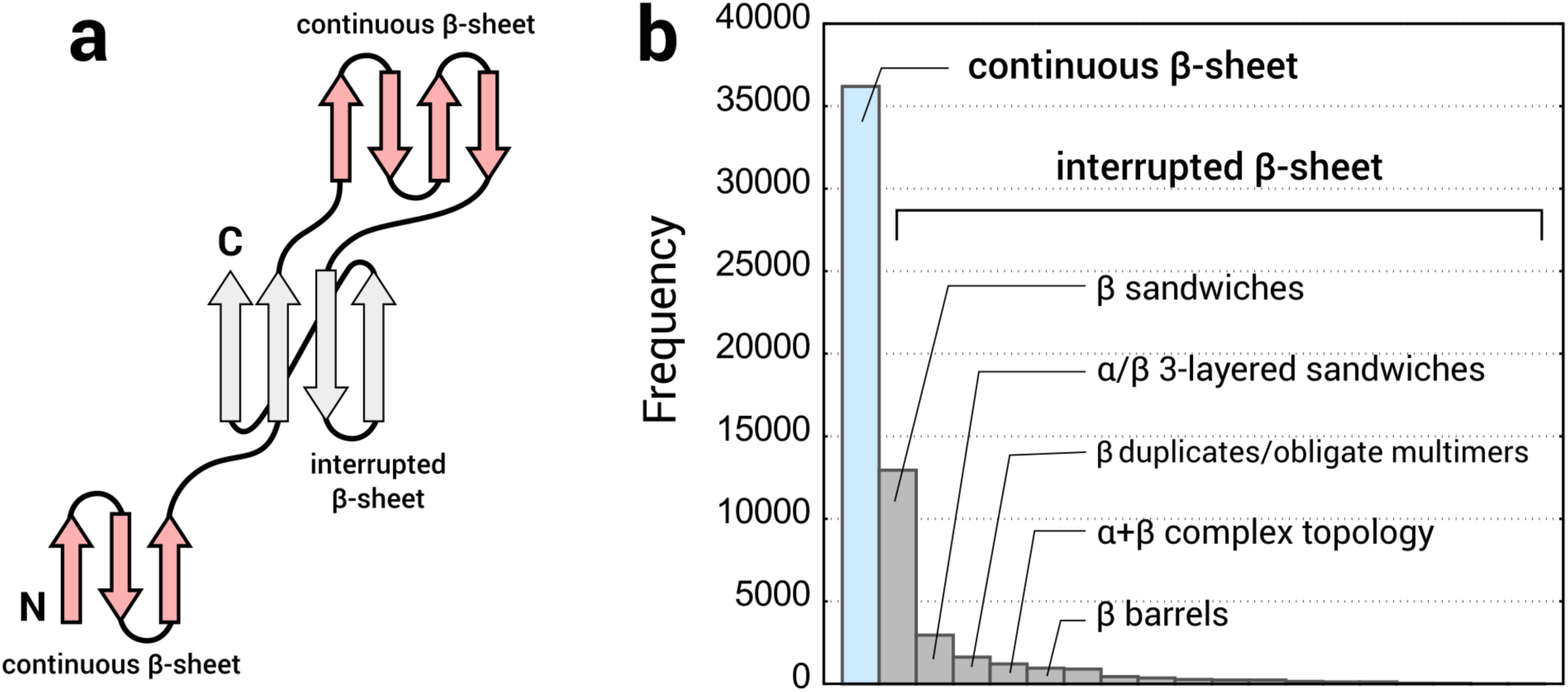
Continuous β-sheet and interrupted β-sheet. **a**, Illustration for a protein chain comprising three domains, each containing a β-sheet. The red-color domains have continuous β-sheets in which there is no insertion of any other β-sheet domains, whereas the gray-color domain has an interrupted β-sheet in which another β-sheet is inserted in the loop region immediately after the first strand. **b**, Observation frequencies for the continuous and interrupted β-sheets in the ECOD dataset. The blue and gray bars indicate the frequencies for the continuous and interrupted β-sheets, respectively. For the interrupted β-sheets, the frequency is further shown depending on each topology pattern (that is, ‘Architecture’ described in ECOD); the most typical one is the β-sandwich type, consisting of the two β-sheets facing each other with an entangled chain (e.g. Immunoglobulin-like fold).

**Extended Data Fig. 8.**
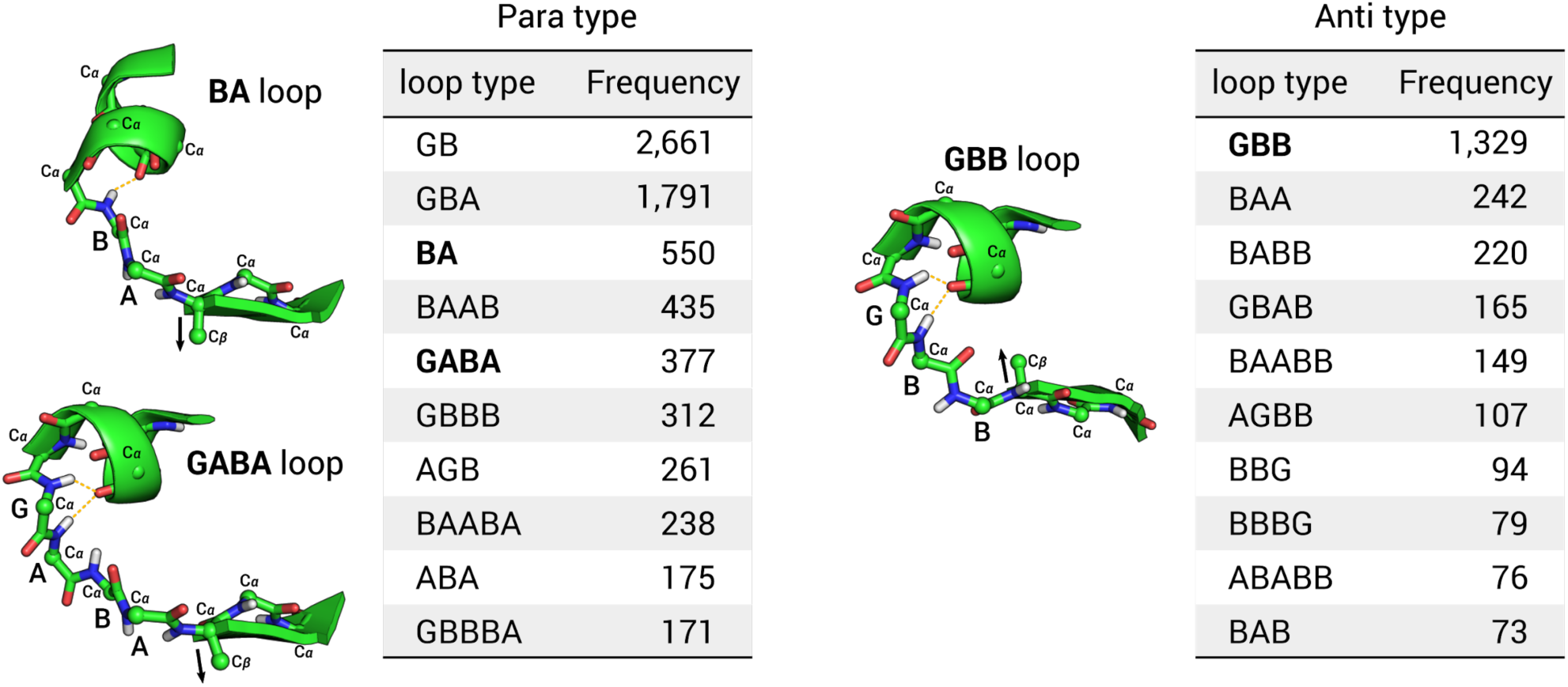
Newly introduced loop patterns for αβ-units. Frequencies of ABEGO torsion patterns for the loops in αβ-units in naturally occurring proteins are shown for the para- and anti-types in the left and right tables, respectively (the para type: the vector from the Cα to Cβ atoms of the first strand residue points away from the helix; the anti type: the same vector points toward the helix). The GB, GBA, and BAAB loops were used in the previous *de novo* designed proteins^9,10^. The BA and GABA loops for the para type and the GBB loop for the anti type were newly introduced in this paper.

**Extended Data Fig. 9.**
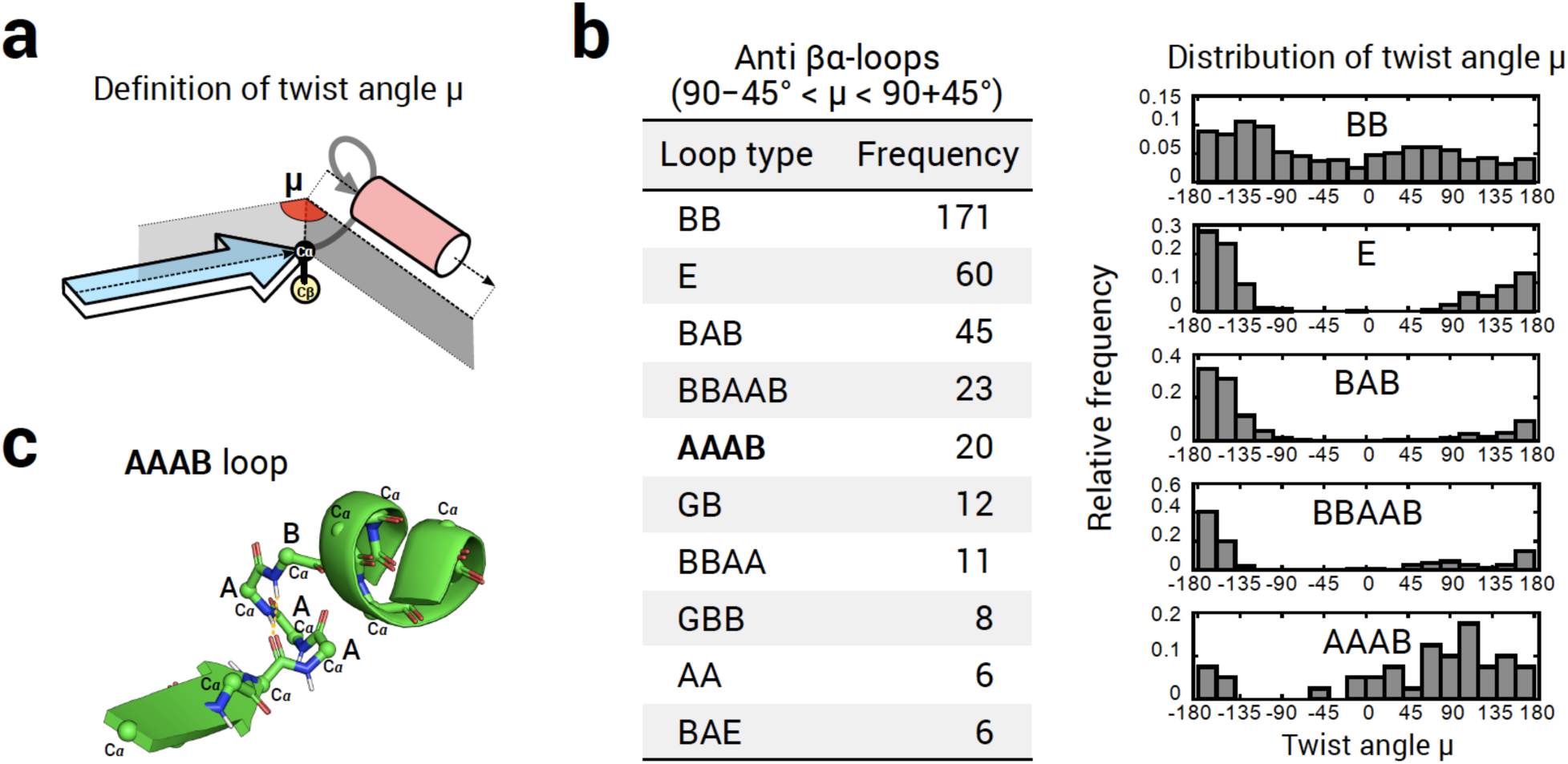
The AAAB loop with the right twist angle for βα-units. **a**, The twist angle μ for βα-units with the anti type^9^ (the vector from the Cα to Cβ atoms of the last strand residue points away from the helix) was defined as the dihedral angle between the plane defined by the β-strand vector and the CαCβ vector of the last strand residue, and the plane defined by the same CαCβ vector and the α-helix vector (the definitions for the β-strand and α-helix vectors are the same as those described in the paper^9^. **b**, Left: the frequencies for ABEGO torsion patterns of loops in βα-units, having the μ angle around 90°, in naturally occurring protein structures. Right: Distributions of the twist angle μ for each of the most frequently observed five loop types in the left table. The AAAB loop showing the clear peak at ∼90° was used in the NF7 fold design. **c**, The backbone structure of the AAAB loop.

## Extended Data Tables

**Extended Data Table 1.** Summary of experimental results for designed proteins.

| | #designs<br>tested | Expressed<sup>1</sup> | Soluble<sup>1</sup> | $\alpha\beta$-protein<br>CD spectrum<br>(20 °C) | Monomeric<sup>2</sup> | Well resolved<br>HSQC<sup>3</sup> | Success<br>(rate %) |
| --- | --- | --- | --- | --- | --- | --- | --- |
| NF1 | 16 | 16 | 14 | 14 | 6 | 4 | 4 (25) |
| NF2 | 4 | 4 | 4 | 4 | 4 | 2 | 2 (50) |
| NF3 | 4 | 4 | 3 | 3 | 3 | 2 | 2 (50) |
| NF4 | 6 | 6 | 6 | 6 | 5 | 4 | 4 (67) |
| NF5 | 6 | 6 | 6 | 6 | 2 | 2 | 2 (33) |
| NF6 | 6 | 6 | 6 | 6 | 5 | 5 | 5 (83) |
| NF7 | 6 | 6 | 6 | 6 | 3 | 3 | 3 (50) |
| NF8 | 12 | 12 | 11 | 11 | 6 | 6 | 6 (50) |

The second column shows the number of designs experimentally tested for the fold in the leftmost column. The subsequent columns give the number of designs that satisfy each experimental characterization, which was performed sequentially from the left to the right. The successful designs are defined as those that satisfy all criteria and are expected to fold into the correct fold. The details of the results are shown in Supplementary Tables 9-16.
<sup>1</sup> Expression and solubility were assessed by SDS-PAGE and mass spectrometry.
<sup>2</sup> SEC-MALS was used to determine oligomerization state. The number of designs in which the main peak of the absorbance at 280 nm corresponds to the monomeric state was counted.
<sup>3</sup> <sup>1</sup>H-<sup>15</sup>N HSQC spectra were collected.

**Extended Data Table 2.** NMR constraints and structure statistics of 8 designed structures.

|  |  |  |  |  |
| --- | --- | --- | --- | --- |
| Design ID | NF1-14 | NF2-02 | NF3-03 | NF4-04 |
| PDB ID | 7BPL | 7BPM | 7BQE | 7BQC |
| BMRB Entry | 36327 | 36328 | 36334 | 36332 |
| **NMR distance constraints** |  |  |  |  |
| **Detected NOE peaks** |  |  |  |  |
| <sup>15</sup>N-edited NOESY | 1,662 | 1,256 | 1,622 | 1,313 |
| <sup>13</sup>C-edited NOESY for aliphatic | 4,012 | 1989 | 2,698 | 3,737 |
| <sup>13</sup>C-edited NOESY for aromatic | 147 | 63 | 135 | 118 |
| **Distance constraints** |  |  |  |  |
| Total | 2597 (100.0%) | 1383 (100.0%) | 2,160 (100.0%) | 1,944 (100.0%) |
| Intra-residue | 525 (20.2%) | 319 (23.1%) | 529 (24.5%) | 493 (25.4%) |
| Inter-residue |  |  |  |  |
| Sequential (\|i-j\| = 1) | 672 (25.9%) | 391 (28.3%) | 565 (26.2%) | 481 (24.7%) |
| Medium-range (1 < \|i-j\| < 5) | 585 (22.5%) | 341 (24.7%) | 498 (23.1%) | 375 (19.3%) |
| Long-range (\|i-j\| ≥ 5) | 816 (31.4%) | 332 (24.0%) | 568 (26.3%) | 595 (30.6%) |
| **Dihedral constraints** |  |  |  |  |
| Total | 167 | 114 | 178 | 200 |
| **NMR structure statistics** |  |  |  |  |
| **Target function (Å<sup>2</sup>)** |  |  |  |  |
| Total | 1.61±0.03 | 0.35±0.008 | 0.66±0.11 | 1.06±0.11 |
| Upl<sup>¶</sup> | 0.0045±0.0004 | 0.0027±0.0002 | 0.0044±0.0019 | 0.0023±0.0006 |
| VDW | 6.7±0.10 | 1.9±0.10 | 3.0±0.50 | 5.0±0.50 |
| Dihedral | 0.69±0.02 | 0.38±0.02 | 0.31±0.03 | 0.56±0.06 |
| **Amber structure statistics (kcal/mol)** |  |  |  |  |
| Amber energy | -4606.9 | -3761.9 | -5040.9 | -5591.5 |
| Restrains distance | 15.42 | 6.46 | 6.805 | 7.095 |
| Restrains dihedral angle | 7.21 | 2.84 | 2.740 | 5.523 |
| **RMSD\* (Å)** |  |  |  |  |
| Backbone (C', N, Cα, O) | 0.207 | 0.207 | 0.403 | 0.449 |
| **RDC evaluation<sup>§</sup>** |  |  |  |  |
| Total number of RDC values | 80 | 52 | 85 | 93 |
| RMS error | 0.904 (0.894) | 0.923 (0.916) | 0.920 (0.862) | 0.917 (0.918) |
| Q-value | 0.338 (0.330) | 0.377 (0.397) | 0.326 (0.378) | 0.380 (0.378) |
| **Ramachandran plot** |  |  |  |  |
| Most favored regions (%) | 95 | 99 | 99 | 99 |
| Allowed regions (%) | 4 | 1 | 1 | 1 |
| Outliers (%) | 2 | 0 | 0 | 0 |
| Design ID | NF5-03 | NF6-02 | NF7-04 | NF8-01 |
| PDB ID | 7BPP | 7BQB | 7BPN | 7BQD |
| BMRB Entry | 36330 | 36331 | 36329 | 36333 |
| **NMR distance constraints** |  |  |  |  |
| **Detected NOE peaks** |  |  |  |  |
| <sup>15</sup>N-edited NOESY | 1,214 | 1,141 | 1,306 | 1,001 |
| <sup>13</sup>C-edited NOESY for aliphatic | 2,773 | 2,347 | 3,259 | 2,527 |
| <sup>13</sup>C-edited NOESY for aromatic | 128 | 115 | 176 | 93 |
| **Distance constraints** |  |  |  |  |
| Total | 1943 (100.0%) | 1802 (100.0%) | 1902 (100.0%) | 1,654 (100.0%) |
| Intra-residue | 502 (25.8%) | 467 (25.9%) | 525 (27.6%) | 402 (24.5%) |
| Inter-residue |  |  |  |  |
| Sequential ($\vert i-j\vert = 1$) | 458 (23.6%) | 457 (25.4%) | 473 (24.9%) | 449 (27.1%) |
| Medium-range ($1 < \vert i-j\vert < 5$) | 380 (19.6%) | 397 (22.0%) | 351 (18.5%) | 326 (19.7%) |
| Long-range ($\vert i-j\vert \geq 5$) | 603 (31.0%) | 481 (26.7%) | 553 (29.1%) | 477 (28.8%) |
| **Dihedral constraints** |  |  |  |  |
| Total | 187 | 183 | 214 | 127 |
| **NMR structure statistics** |  |  |  |  |
| **Target function ($\text{\AA}^2$)** |  |  |  |  |
| Total | $0.65 \pm 0.05$ | $0.57 \pm 0.02$ | $0.86 \pm 0.02$ | $0.57 \pm 0.02$ |
| Upl<sup>¶</sup> | $0.0022 \pm 0.0006$ | $0.0022 \pm 0.0002$ | $0.0015 \pm 0.0005$ | $0.0025 \pm 0.0006$ |
| VDW | $3.4 \pm 0.30$ | $2.3 \pm 0.10$ | $3.2 \pm 0.20$ | $2.4 \pm 0.20$ |
| Dihedral | $0.49 \pm 0.06$ | $0.31 \pm 0.07$ | $0.53 \pm 0.04$ | $0.38 \pm 0.02$ |
| **Amber structure statistics (kcal/mol)** |  |  |  |  |
| Amber energy | -5256.5 | -5041.9 | -5404.4 | -3731.3 |
| Restraints distance | 8.295 | 6.865 | 5.817 | 5.926 |
| Restraints dihedral angle | 6.275 | 2.740 | 8.527 | 3.725 |
| **RMSD\* ($\text{\AA}$)** |  |  |  |  |
| Backbone (C', N, Cα, O) | 0.360 | 0.424 | 0.477 | 0.332 |
| **RDC evaluation<sup>§</sup>** |  |  |  |  |
| Total number of RDC values | 89 | 78 | 98 | 57 |
| RMS error | 0.909 (0.937) | 0.955 (0.924) | 0.915 (0.893) | 0.930 (0.926) |
| Q-value | 0.411 (0.340) | 0.319 (0.403) | 0.404 (0.458) | 0.317 (0.329) |
| **Ramachandran plot** |  |  |  |  |
| Most favored regions (%) | 100 | 100 | 98 | 99 |
| Allowed regions (%) | 0 | 0 | 2 | 1 |
| Outliers (%) | 0 | 0 | 0 | 0 |

<sup>¶</sup> The Upl target function becomes more than zero when atom pairs are beyond the upper limits of distance constraints.
\* The RMSD values were calculated for the 20 structures overlaid to the mean coordinates for the ordered regions, which were automatically identified by Fit\_Robot using multi-dimensional non-linear scaling<sup>60</sup>.
<sup>§</sup> The RDC back-calculation was performed by PALES<sup>61</sup> using experimentally determined values of 1-bond <sup>1</sup>H-<sup>15</sup>N RDC. The averaged correlation RMS and Q-value for NMR structure ensemble and design coordinates (as shown in parentheses) between the simulated and experimental values was obtained using the <sup>1</sup>H-<sup>15</sup>N signals except completely overlapped peaks or the residues in low order-parameters (less than 0.8) predicted by TALOS+.

## References

1. Orengo, C. A., Jones, D. T. & Thornton, J. M. Protein superfamilles and domain superfolds. Nature 372, 631 (1994).

2. Murzin, A. G., Brenner, S. E., Hubbard, T. & Chothia, C. SCOP: a structural classification of proteins database for the investigation of sequences and structures. J. Mol. Biol. 247, 536–540 (1995).

3. Orengo, C. A. et al. CATH–a hierarchic classification of protein domain structures. Structure 5, 1093–1109 (1997).

4. Zhang, Y., Hubner, I. A., Arakaki, A. K., Shakhnovich, E. & Skolnick, J. On the origin and highly likely completeness of single-domain protein structures. Proceedings of the National Academy of Sciences 103, 2605–2610 (2006).

5. Taylor, W. R., Chelliah, V., Hollup, S. M., MacDonald, J. T. & Jonassen, I. Probing the ‘dark matter’ of protein fold space. Structure 17, 1244–1252 (2009).

6. Cossio, P. et al. Exploring the universe of protein structures beyond the Protein Data Bank. PLoS Comput. Biol. 6, e1000957 (2010).

7. Chitturi, B., Shi, S., Kinch, L. N. & Grishin, N. V. Compact structure patterns in proteins. J. Mol. Biol. 428, 4392–4412 (2016).

8. Kuhlman, B. et al. Design of a novel globular protein fold with atomic-level accuracy. Science 302, 1364–1368 (2003).

9. Koga, N. et al. Principles for designing ideal protein structures. Nature 491, 222 (2012).

10. Lin, Y.-R. et al. Control over overall shape and size in de novo designed proteins. Proceedings of the National Academy of Sciences 112, E5478–E5485 (2015).

11. Huang, P.-S. et al. De novo design of a four-fold symmetric TIM-barrel protein with atomic-level accuracy. Nat. Chem. Biol. 12, 29 (2016).

12. Marcos, E. et al. Principles for designing proteins with cavities formed by curved β sheets. Science 355, 201–206 (2017).

13. Marcos, E. et al. De novo design of a non-local β-sheet protein with high stability and accuracy. Nat. Struct. Mol. Biol. 25, 1028 (2018).

14. Dou, J. et al. De novo design of a fluorescence-activating β-barrel. Nature 561, 485 (2018).

15. Richardson, J. S. Handedness of crossover connections in beta sheets. Proceedings of the National Academy of Sciences 73, 2619–2623 (1976).

16. Richardson, J. S. β-Sheet topology and the relatedness of proteins. Nature 268, 495 (1977).

17. Grainger, B., Sadowski, M. I. & Taylor, W. R. Re-evaluating the ‘rules’ of protein topology. J. Comput. Biol. 17, 1371–1384 (2010).

18. Martin, A. C. R., et al. Protein folds and functions. Structure 6, 875–884 (1998).

19. Orengo, C. A. et al. The CATH Database provides insights into protein structure/function relationships. Nucleic Acids Res. 27, 275–279 (1999).

20. Zhang, C. & Kim, S.-H. The anatomy of protein β-sheet topology. J. Mol. Biol. 299, 1075–1089 (2000).

21. Ruczinski, I., Kooperberg, C., Bonneau, R. & Baker, D. Distributions of beta sheets in proteins with application to structure prediction. Proteins: Struct. Funct. Bioinf. 48, 85–97 (2002).

22. Wintjens, R. T., Rooman, M. J. & Wodak, S. J. Automatic classification and analysis of αα-turn motifs in proteins. J. Mol. Biol. 255, 235–253 (1996).

23. Rohl, C. A., Strauss, C. E. M., Misura, K. M. S. & Baker, D. Protein structure prediction using Rosetta. in Methods in enzymology vol. 383 66–93 (Elsevier, 2004).

24. Bradley, P., Misura, K. M. S. & Baker, D. Toward high-resolution de novo structure prediction for small proteins. Science 309, 1868–1871 (2005).

25. Zhang, Y. & Skolnick, J. TM-align: a protein structure alignment algorithm based on the TM-score. Nucleic Acids Res. 33, 2302–2309 (2005).

26. Minami, S., Sawada, K. & Chikenji, G. MICAN: a protein structure alignment algorithm that can handle Multiple-chains, Inverse alignments, C α only models, Alternative alignments, and Non-sequential alignments. BMC Bioinformatics 14, 24 (2013).

27. Minami, S., Sawada, K., Ota, M. & Chikenji, G. MICAN-SQ: a sequential protein structure alignment program that is applicable to monomers and all types of oligomers. Bioinformatics 34, 3324–3331 (2018).

28. Gilbert, D., Westhead, D., Nagano, N. & Thornton, J. Motif-based searching in TOPS protein topology databases. Bioinformatics 15, 317–326 (1999).

29. Leaver-Fay, A. et al. ROSETTA3: an object-oriented software suite for the simulation and design of macromolecules. Methods Enzymol. 487, 545–574 (2011).

30. Dou, J. et al. De novo design of a fluorescence-activating β-barrel. Nature 561, 485–491 (2018).

31. Kobayashi, N. et al. KUJIRA, a package of integrated modules for systematic and interactive analysis of NMR data directed to high-throughput NMR structure studies. J. Biomol. NMR 39, 31–52 (2007).

32. Kobayashi, N. et al. Noise peak filtering in multi-dimensional NMR spectra using convolutional neural networks. Bioinformatics 34, 4300–4301 (2018).

33. Marcos, E. et al. De novo design of a non-local β-sheet protein with high stability and accuracy. Nat. Struct. Mol. Biol. 25, 1028–1034 (2018).

34. Burton, A. J., Thomson, A. R., Dawson, W. M., Brady, R. L. & Woolfson, D. N. Installing hydrolytic activity into a completely de novo protein framework. Nat. Chem. 8, 837–844 (2016).

35. Chevalier, A. et al. Massively parallel de novo protein design for targeted therapeutics. Nature 550, 74 (2017).

36. Thomas, F. et al. De Novo-Designed α-Helical Barrels as Receptors for Small Molecules. ACS Synth. Biol. 7, 1808–1816 (2018).

37. Langan, R. A. et al. De novo design of bioactive protein switches. Nature 572, 205–210 (2019).

38. Silva, D.-A. et al. De novo design of potent and selective mimics of IL-2 and IL-15. Nature 565, 186 (2019).

39. Chen, Z. et al. De novo design of protein logic gates. Science 368, 78–84 (2020).

40. Linsky, T. W. et al. De novo design of potent and resilient hACE2 decoys to neutralize SARS-CoV-2. Science 370, 1208–1214 (2020).

41. Vorobieva, A. A. et al. De novo design of transmembrane β barrels. Science 371, (2021).

42. Quijano-Rubio, A. et al. De novo design of modular and tunable protein biosensors. Nature (2021) doi:10.1038/s41586-021-03258-z.

43. Sesterhenn, F. et al. De novo protein design enables the precise induction of RSV-neutralizing antibodies. Science 368, (2020).

44. Cao, L. et al. De novo design of picomolar SARS-CoV-2 miniprotein inhibitors. Science 370, 426–431 (2020).

45. Wang, G. & Dunbrack, R. L., Jr. PISCES: a protein sequence culling server. Bioinformatics 19, 1589–1591 (2003).

46. Cheng, H. et al. ECOD: an evolutionary classification of protein domains. PLoS Comput. Biol. 10, e1003926 (2014).

47. Xu, D. & Zhang, Y. Improving the physical realism and structural accuracy of protein models by a two-step atomic-level energy minimization. Biophys. J. 101, 2525–2534 (2011).

48. Frishman, D. & Argos, P. Knowledge-based protein secondary structure assignment. Proteins: Struct. Funct. Bioinf. 23, 566–579 (1995).

49. O’Meara, M. J. et al. Combined covalent-electrostatic model of hydrogen bonding improves structure prediction with Rosetta. J. Chem. Theory Comput. 11, 609–622 (2015).

50. Alford, R. F. et al. The Rosetta All-Atom Energy Function for Macromolecular Modeling and Design. J. Chem. Theory Comput. 13, 3031–3048 (2017).

51. Kuhlman, B. & Baker, D. Native protein sequences are close to optimal for their structures. Proceedings of the National Academy of Sciences 97, 10383–10388 (2000).

52. Sheffler, W. & Baker, D. RosettaHoles: rapid assessment of protein core packing for structure prediction, refinement, design, and validation. Protein Sci. 18, 229–239 (2009).

53. Jones, D. T. Protein secondary structure prediction based on position-specific scoring matrices. J. Mol. Biol. 292, 195–202 (1999).

54. Jansson, M. et al. High-level production of uniformly 15 N-and 13 C-enriched fusion proteins in Escherichia coli. J. Biomol. NMR 7, 131–141 (1996).

55. Pace, C. N., Vajdos, F., Fee, L., Grimsley, G. & Gray, T. How to measure and predict the molar absorption coefficient of a protein. Protein Sci. 4, 2411–2423 (1995).

56. Schanda, P., Van Melckebeke, H. & Brutscher, B. Speeding up three-dimensional protein NMR experiments to a few minutes. J. Am. Chem. Soc. 128, 9042–9043 (2006).

57. Schmidt, E. & Güntert, P. A new algorithm for reliable and general NMR resonance assignment. J. Am. Chem. Soc. 134, 12817–12829 (2012).

58. Shen, Y., Delaglio, F., Cornilescu, G. & Bax, A. TALOS+: a hybrid method for predicting protein backbone torsion angles from NMR chemical shifts. J. Biomol. NMR 44, 213–223 (2009).

59. Güntert, P. & Buchner, L. Combined automated NOE assignment and structure calculation with CYANA. J. Biomol. NMR 62, 453–471 (2015).

60. Kobayashi, N. A robust method for quantitative identification of ordered cores in an ensemble of biomolecular structures by non-linear multi-dimensional scaling using inter-atomic distance variance matrix. J. Biomol. NMR 58, 61–67 (2014).

61. Zweckstetter, M. & Bax, A. Prediction of Sterically Induced Alignment in a Dilute Liquid Crystalline Phase: Aid to Protein Structure Determination by NMR. J. Am. Chem. Soc. 122, 3791–3792 (2000).

62. Finkelstein, A. V. & Ptitsyn, O. B. Why do globular proteins fit the limited set of foldin patterns? Prog. Biophys. Mol. Biol. 50, 171–190 (1987).

63. Jamroz, M. et al. KnotProt: a database of proteins with knots and slipknots. Nucleic Acids Res. 43, D306–D314 (2014).

