## Supplementary FIle for "Exploration of novel αβ-protein folds through *de novo* design"

### Supplementary methods

#### NMR structure determination

All αβ-proteins in this study are generally stable and have long lifetime (more than 2~3 weeks) at NMR concentration (0.5~1.0mM), and rarely minor components can be found. On the other hand, there is no noticeable signal change in 2D $^{1}H$-$^{15}N$ HSQC at different concentrations (2~10 times dilution) for all samples. Taken together with the results from SEC-MALS analysis, all the samples are considered to be in stable monomeric states throughout the NMR experiments.

Prior to the structure determination, the analyst never has known the designed structure even its sequence (perfectly blind analysis) in order to avoid any arbitrary bias for the automated NMR analysis. Owing to the high sensitivity of modern NMR machines (700~800 MHz equipped with 2$^{nd}$ or 3$^{rd}$ generation of Cryo-probe) and high concentration of samples, significant number of NOE peaks were yielded from NOESY type spectra. More than 80~90% of NOE peaks are assigned by CYANA to obtain well converged structure, supporting high consensus between NOE peaks and calculated NMR structure.

Several CYANA calculations were performed using the Acs table, NOE peak table and dihedral angle constraints to obtain 20 models with lowest target functions. For the obtained CYANA structures, implicit water refinement calculations were performed by AMBER12 with ff99SB force field. The dihedral angle constraints and distance constraints including additional chirality and backbone omega angle constraints were converted for AMBER format using SANDER tool. In each initial stage of the refinement, energy minimization of 500 steps (250 step: steepest gradient, followed by 250 step: conjugate gradient decent) without electrostatic energy and NMR constraint terms was carried out. A short molecular dynamics calculation (total 30 psec, time step 1.0 fsec, using SHAKE algorism) was followed using electrostatic energy based on generalized Born model (salt concentration: 0.1 M, disabled Surface Accessibility (SA) function, electrostatic potential radius cutoff: 18 Å) and NMR constraint terms. The temperature was gradually increased from 0 K to 300 K for 1,500 steps, then run dynamic calculation at 300 K for 1,500 steps. In the final stage of the refinement, 2,000 steps of energy minimization was performed with the same energy terms.

The RDC back calculations were used for validating the determined NMR structures. This strategy enhances the reliability of the determined NMR structure, with the geometrical normality (such as Ramachandran plots and VdW clash) and violations for restraints. Although residual dipolar couplings (RDC) can be affected by local motion of HN-N vector in wide-range of time scale, the effect is not big if a sufficient number (coverage of residues more than 80%) and sufficient amplitude of RDC values (>±10Hz) are obtained. Using software PALES (*1*), each model coordinates of calculated NMR ensemble and a number of experimental RDC values, by means of singular value decomposition (SVD), Saupe matrix can be obtained to estimate Euler angles and amplitudes of alignment tensor. Then using the tensor parameter, PALES can calculate RMS error between simulated and experimental RDC values. The RMS error greater than 0.9 means the calculated structure can be trustful, unless the RDC values were used for structure calculation as constraints. For the severely overlapped residues and the residues with low order parameter (less than 0.8 predicted by TALOS+), the RDC values were eliminated from the analysis. It would be noteworthy that more than 80% of observed RDC data were used for all of the RDC analysis in this study. Fortunately, the designed αβ-proteins in this study composed of a few helices and slightly twisting β-sheets which are largely different orientation, so the RDC analysis using $^{1}D_{1H\text{-}15N}$ is firmly suitable for structure validation.

NF1-14

The methyl protons of Ile46-Hγ2, Ile28-Hδ1, Leu103-Hδ1/2 are weakly shielded by the aromatic rings of Phe80 and Trp99 respectively. Interestingly side-chain of Gln18 is stacked by Tyr38, which is well consistent with the strongly shielded amide proton of side-chain Gln18. Unusually down-filed shifted Gln13-Hε2 indicates formation of strong hydrogen bond stabilizing and tightly packing between helix1 and helix3. All of the aromatic rings are nearly the same location when overlaying designed and NMR structures, which supports both of the structures have strikingly same structure. As shown by 2D $^{1}H$-$^{15}N$ HSQC in Supplementary Figure 1, NF1 does not have any minor components at the NMR condition. In the RDC validation analysis, NMR structure was slightly better than designed one (RMS were 0.904 and 0.894, respectively). This fact strongly supports NMR and designed structures are completely same structure in solution.

NF2-02

$^{1}H$-$^{15}N$ HSQC is shown in Supplementary Figure 2. There are two tyrosine in this protein. Ile27-Hδ1 and -Hγ2 methyl protons are shielded by aromatic ring of Tyr16. Gln53-Hε1/2 are slightly shielded, suggesting ring current effect by Tyr63. As a result of the RDC validation analysis, RMS error is slightly better than designed structure (0.927 and 0.916, respectively). Taken together with the shielded protons and RDC scores, the protein in solution has designed structure demonstrated by NMR analysis.

NF3-03

$^{1}H$-$^{15}N$ HSQC is shown in Supplementary Figure 3. The methyl groups, Ile49-Hγ2, -Hγ1, Ile71-Hδ1, -Hγ2 and Hγ1, are shielded by Phe54, while despite Ile96 close to Trp101, no protons are shielded. Val32 is also close enough to Tyr26 but no shielded proton was found. Although designed structure does not contain Trp101, a lot of NOE from ring to methyl protons were observed. Since any of the proximate methyls are not shielded, Trp101 may cover the hydrophobic cluster of the methyl groups but not in a fixed orientation. This would result in cancelation of ring current effect to the methyl protons.

Instead of these facts, Arg24-Hβ2/3 are obviously shielded, weakly stacking to Tyr26. This interaction is not found in designed structure. The key residues Phe54 and Tyr26 are the same location and orientation which well explain shielding effect to the methyl and methylene protons. In the overlaid designed and NMR structures, helix4 is slightly away from the hydrophobic core, which may be caused by less number of long-range distance constraints from the helix. The RDC validation analysis showed NMR structure is better than designed one. As the RMS error is greater than 0.9, the solution structure is trustful and similar to the designed structure.

NF4-04

Because of the relatively high pH and salt concentration, a lot of signals which may be located on hairpin are weak, broad and missing in the 2D $^{1}H$-$^{15}N$ HSQC (Supplementary Figure 4). Additionally, there are a lot of aliphatic amino acids such as Ile, Val and Leu (total 33 residues) for this size of protein, so supportive spectra were needed such as 3D (H)CC(CO)NH and (H)CCH-TOCSY for confirmation of sequential assignment and side-chain assignments, and

3D $^{13}$C-HSQC ($^{13}$C-time domain) –NOESY $^{15}$N-HSQC, $^{13}$C-HSQC ($^{13}$C- time domain) NOESY $^{13}$C-HSQC for obtaining methyl-methyl NOEs.

A lot of methyl protons are shielded such as Ile25-Hδ1 by Tyr13, Ile12-Hγ1, -Hδ1 by Phe62, Ile80-Hγ2 and Leu84-Hδ1/2, Val95-Hγ1/2 by Phe28, Leu109-Hδ1/2 by Phe31. Trp116 is not involved in rigid regions, may be in flexible conformation. The side-chain amide signals of Gln88 are very unusual position in 2D $^{1}$H-$^{15}$N HSQC (Supplementary Figure 4), probably tightly packed in the core of protein and forming hydrogen bond to Leu84-CO. The C-terminal helix is slightly away from the core of protein compared with designed structure. The less number of NOEs can be found around the helix, which means the packing capability between C-terminal helix may be relatively weak. Overall conformation of NMR structure is similar to designed one, however, the largest difference was orientation of Phe31 and Leu106. Interestingly the RDC validation score of the NMR and designed structures were greater than 0.9 (0.917 and 0.918), indicating global fold is moderately correct. The side-chain location and orientation in the hydrophobic core except for the above mentioned residues are nearly the same.

NF5-03

In this protein, a lot of shielded methyl protons can be found in 2D $^{1}$H-$^{13}$C HSQC (Supplementary Figure 5). Ile46-Hγ2 and Hδ1, Ile52-Hδ1, -Hγ2 and Ile60-Hγ2 are shielded by ring current of Phe64, and Ile24-Hγ2, by Tyr35. Ile24-Hγ2 and Val90-Hγ1/2 are also shielded by ring current of Phe73. These facts indicate NMR structure has well consensus with chemical shift data. The location and orientation of aromatic rings, in the superimposed design and NMR structures, are slightly different, however, the backbone confirmations are quite similar. RDC validation scores for both designed and NMR structures are greater than 0.90, demonstrating that solution NMR structures are trustful. Despite the RMS score of designed structure is a little better than NMR one, the shielded methyl protons and location and directions of aromatic rings can be explained more reasonable in NMR structure.

NF6-02

Ile15-Hγ2, Leu19-Hδ2 or –Hδ1 are shielded by Tyr4, Leu71-Hδ1/2 are weakly shielded by Tyr46. Interestingly, the side-chain amide signals of Gln25 are broad and unusual position in $^{1}$H-$^{15}$N HSQC (Supplementary Figure 6). This fact can be explained by the formation of hydrogen bonding to backbone carbonyl group of Gly51, stabilizing packing between helix2 and strand2. In the overlaid designed and NMR structures, all of the secondary structures as well as aromatics rings are very similar positions. RDC analysis demonstrated that NMR structure is slightly better than designed structure (0.955 and 0.924, respectively), indicating designed structure can be very similar conformation in solution.

NF7-04

$^{1}$H-$^{15}$N HSQC is shown in Supplementary Figure 7. The methyl group of Leu49-Hδ1/2 are shielded by Tyr92, while Val34-Hγ1 or Hγ2 seems to be shielded by Phe107. Leu18-Hδ1/2 are shielded by Phe6. Tyr92 is shielding Leu49-Hδ1/2 as well as Gly1-Hα2/3. Since this protein has more aromatic residues than the other proteins, the wide variety of shielding effects can be used to indicate how correct the NMR structure is. Interestingly Gln2 and Gln4 are stabilizing the strand1 by stacking their side-chains to hydrophobic core. Aromatic rings except for Tyr5 are similar locations and orientations in both designed and NMR structures. RDC errors are near 0.9 for both designed and NMR structures (0.915 and 0.893, respectively), indicating NMR structure is correctly determined and quite similar to the designed structure.

NF8-01

Methyl protons are shielded by ring current effect such as Ile39-Hδ1 by Tyr27, Leu17-Hδ1/2 and Leu61-Hδ1/2 by Phe6, Leu-Hδ1/2 by Tyr36, indicating the NMR structure is well consistent with chemical shift data as well as all aromatic tightly packed methyl groups enhancing hydrophobic interaction in the core of protein. Interestingly side-chain amide group of Gln65 is nearly involved in hydrophobic core, as revealed from unusual chemical shifts in 2D $^{1}$H-$^{15}$N HSQC (Supplementary Figure 8). The side-chain orientations and locations of hydrophobic residues are very similar in both designed and NMR structures. RDC validation analysis demonstrated very good score (0.930 and 0.926 for NMR and designed structures), indicating the protein folds in solution as exactly designed.

### Supplementary Figures

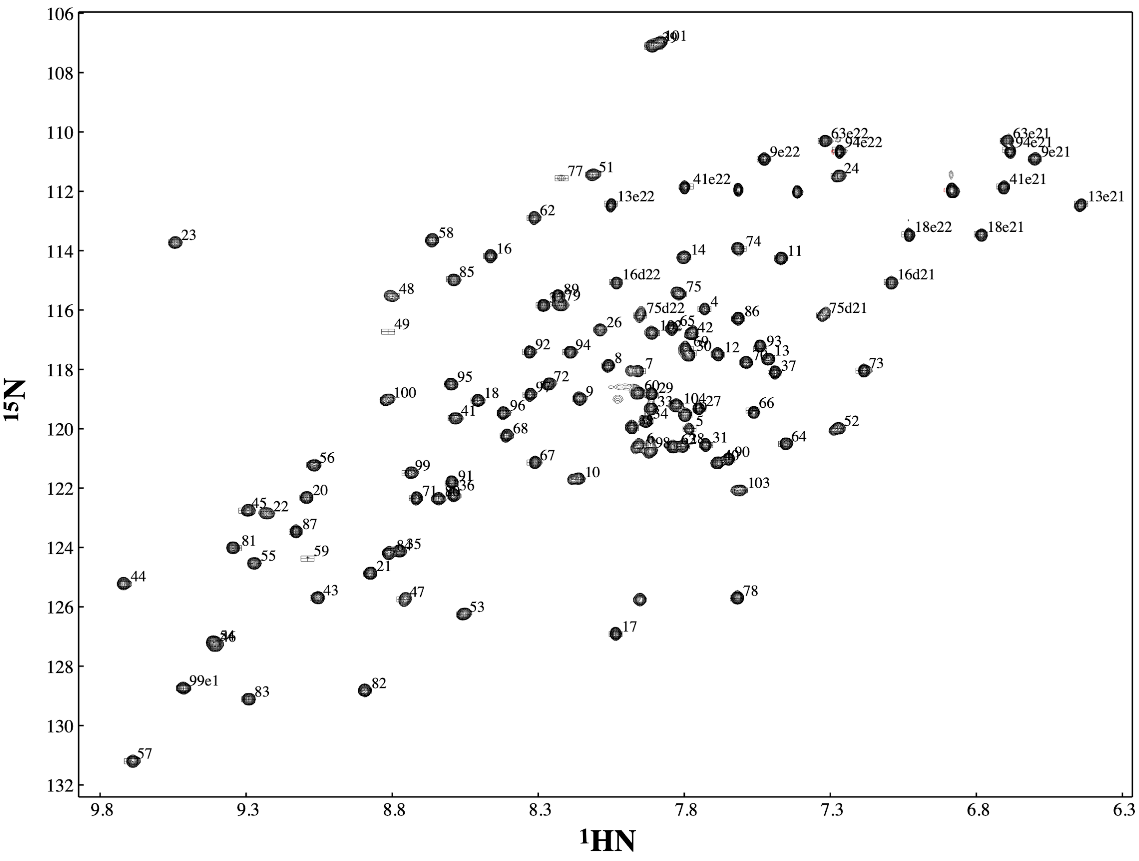


**Supplementary Figure 1 | 2D $^{1}$H-$^{15}$N HSQC of protein NF1-14.**

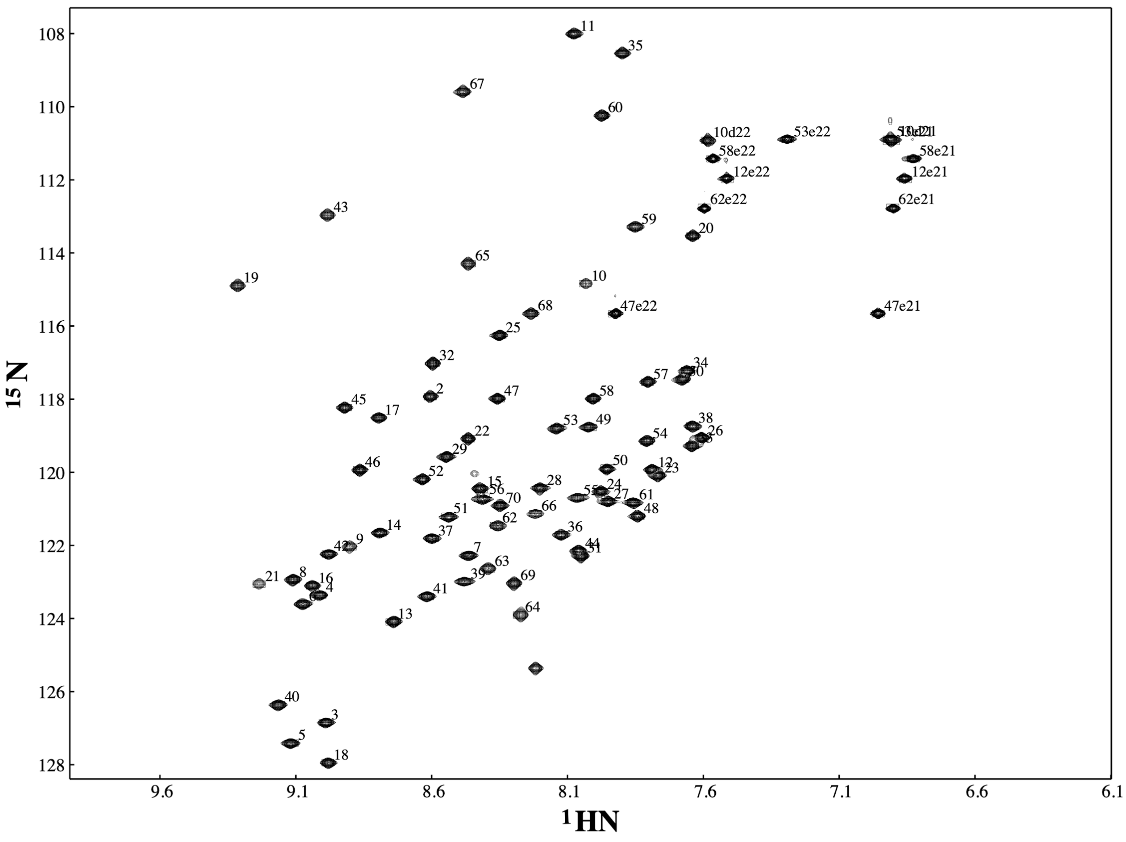


**Supplementary Figure 2 | 2D $^{1}$H-$^{15}$N HSQC of protein NF2-02.**

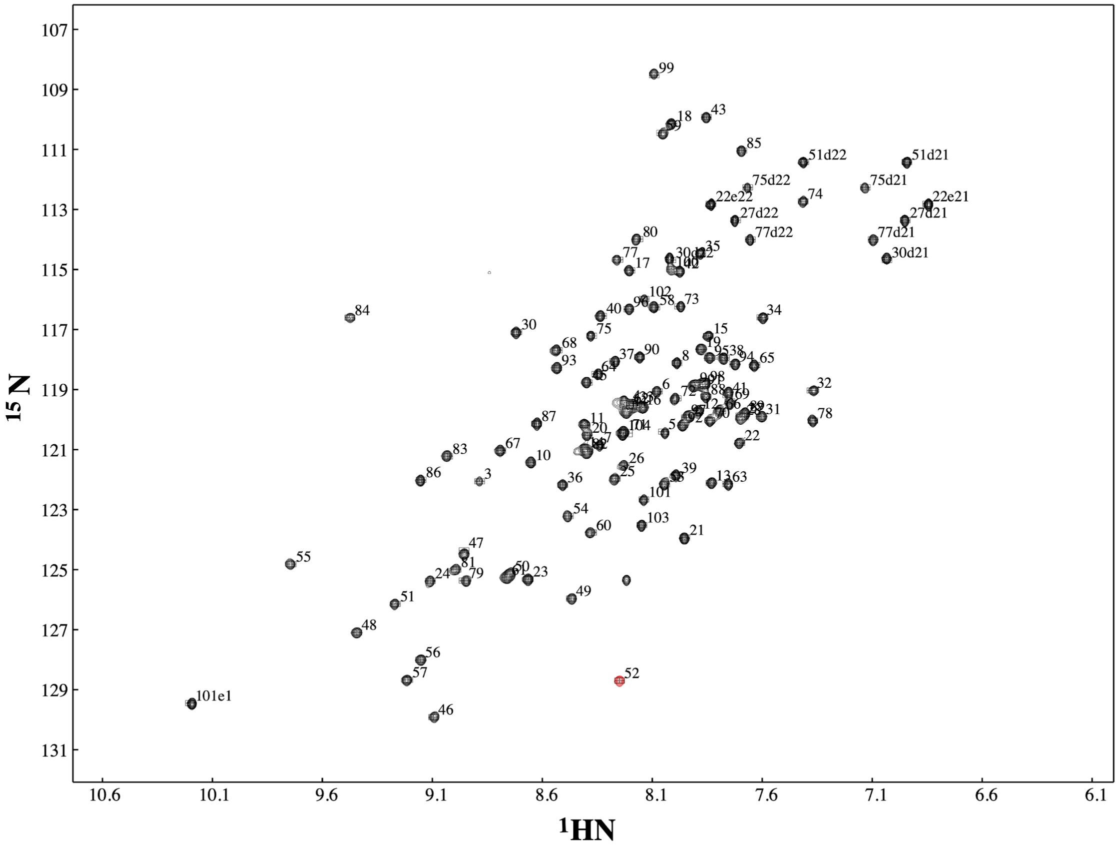


**Supplementary Figure 3 | 2D $^{1}$H-$^{15}$N HSQC of protein NF3-03.**

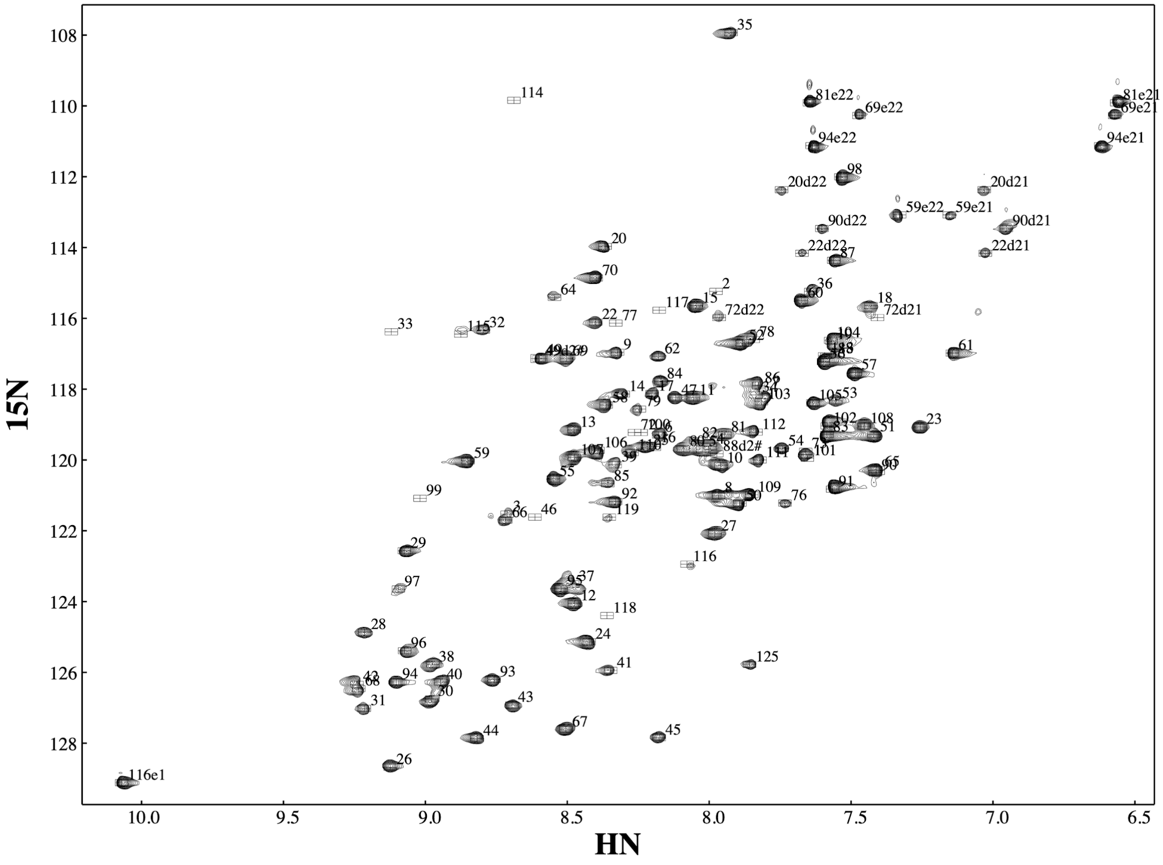


**Supplementary Figure 4 | 2D $^{1}H$-$^{15}N$ HSQC of protein NF4-04.**

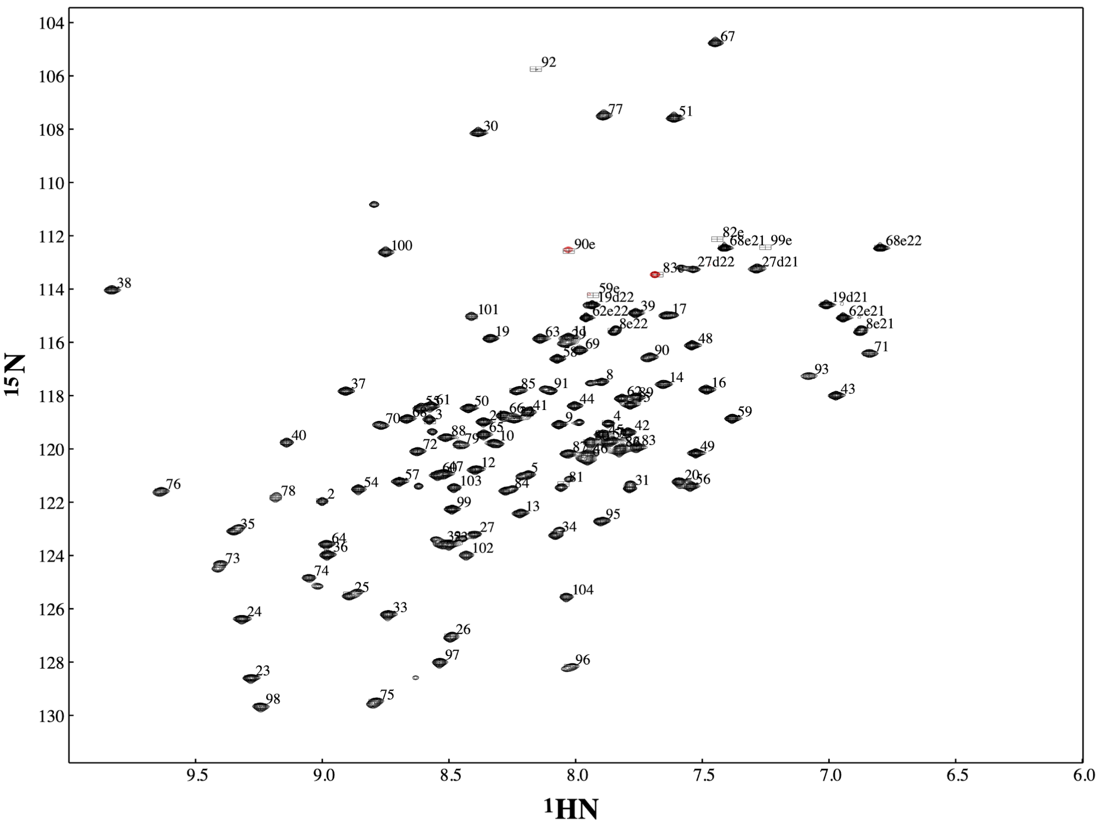


**Supplementary Figure 5 | 2D $^{1}$H-$^{15}$N HSQC of protein NF5-03.**

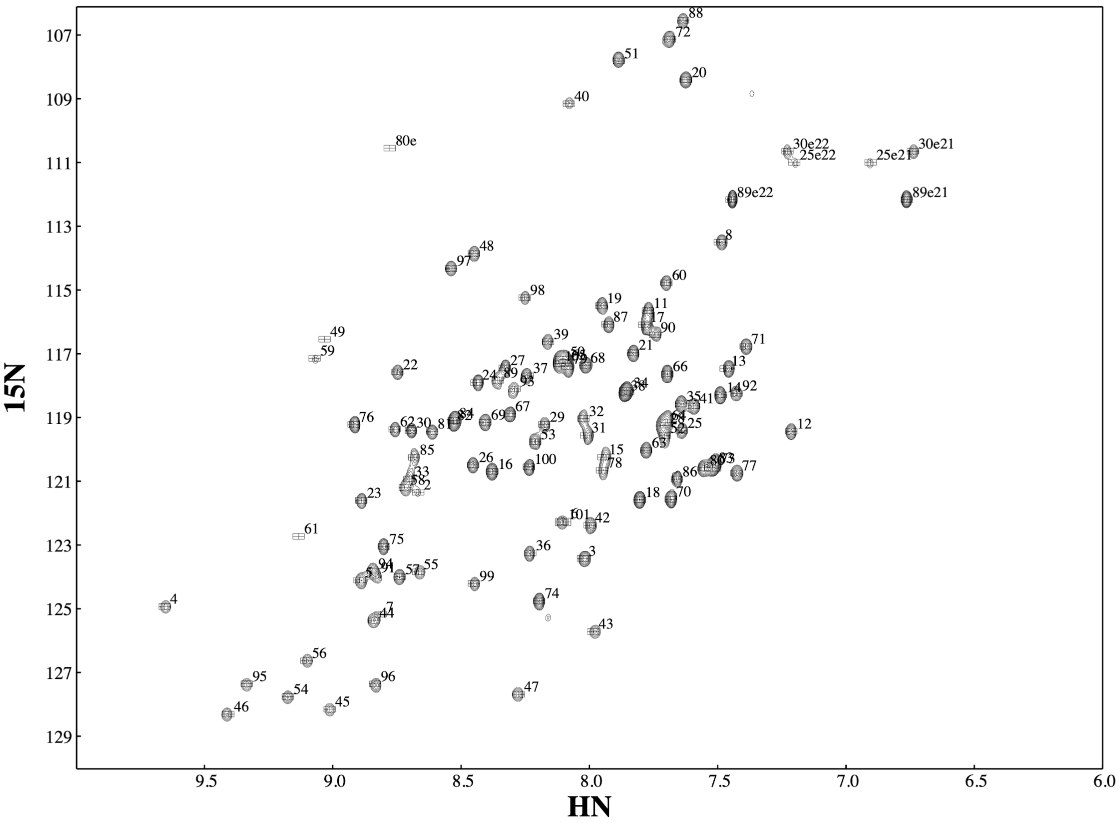


**Supplementary Figure 6 | 2D $^{1}H$-$^{15}N$ HSQC of protein NF6-02.**

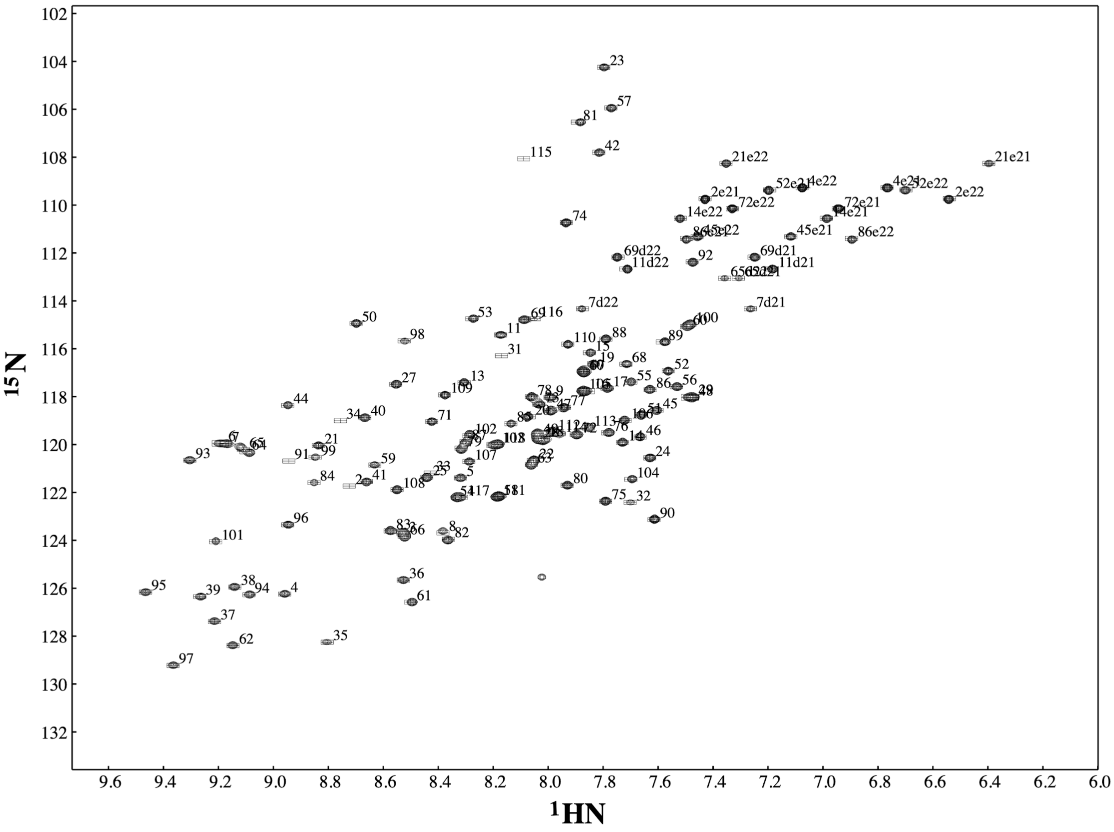


**Supplementary Figure 7 | 2D $^{1}$H-$^{15}$N HSQC of protein NF7-04.**

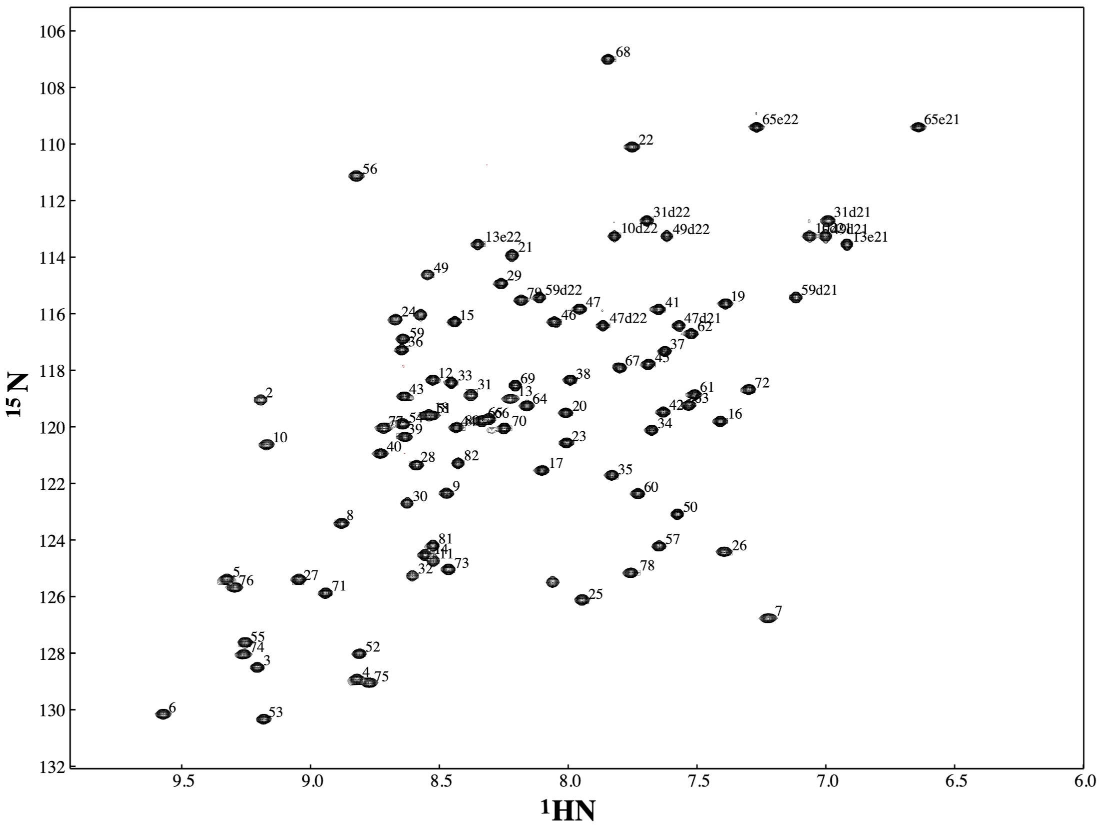


**Supplementary Figure 8 | 2D $^{1}H$-$^{15}N$ HSQC of protein NF8-01.**

### Supplementary Tables

| ID | sequence | E-value |
|---|---|---|
| NF1-01 | mGDLEKIKEAVKRVLPKAKIYEVTSEEEIERVSREIKKEGPRRILVIRSDSGSILIVISRDEKKLR<br>QMSKAAKEVSPNMTLLEFRGQDPEKIRKELERLRRGslehhhhhh | 0.84 |
| NF1-02 | mGDLDKMEEAVKKVNPGAKTYKVTSPEEIKKVSEEIRKEGPRRIIVVRTDSGKILIIISRNEED<br>MRRMSEAIKRVNPNATILEFTGQDPEKIRQRLEELWRGslehhhhhh | 0.14 |
| NF1-03 | mGKLEKMKRAAKKVDPRVKILEVTSEDEIKEASKRIKEEGPRRIVVVRMSQGHILIFISRDEE<br>NMRKIAKAVKKVGPEATILEFEGQDPDRIREELRKLREGswslehhhhhh | 0.55 |
| NF1-04 | mGNLEKIREAVRKVLPKARTLRVNSREDIERASKEIKKEGPRRILIIRSSSGHILIVISRSEEKLE<br>RMKQAAKKVEPDATLLEFRGQDPEKIEKEMRKLYEGslehhhhhh | 3.3 |
| NF1-05 | mGELEKMKRAAKKVDPRVKILEVRTPDQIEEASERIKREGPRRILVVRHDSGNILIFISRDEES<br>LKKIKEAVQRVGPRATTLEFRGQDPEKIERELRKLLKGswslehhhhhh | 0.72 |
| NF1-06 | mGPLDKMAKAAKKVLPKARILRVRTPEQIERASREIKKEGPRRIIIIRADSGNILIIISRDEESAK<br>RIQEAIKRVLPNATLLEFRGQDPEKIEKELRKLLKGswslehhhhhh | 0.007 |
| NF1-07 | mGDVEDIMNDAKNLSPSIQIYEVTTPDELEQAVRDIKNTGAQVVILVWTSSGKLLLFAVTNPE<br>DAERVARDAKKRNPSAKVVRLEGVPPDDIEEQARRLWKGslehhhhhh | 1.2 |
| NF1-08 | mGDVDNVKKRVKEQDPNAQVYEATTPDEIEEVVKRVKKYGAQVVILVYLSDGKIIVIAVRDP<br>SVADQIIEDLKKQNPNLTIIRLEGAPPDELKRRMEELWRGslehhhhhh | 0.21 |
| NF1-09 | mGTVDDIIKDAQNLSPSVQVYEVSTPDEVKEAVERVKNTGAQVVILVYTSSGKLFLFAITDPEI<br>ARKIAEDAKKRNPNAKVKRLEGKPPEDIRQQMEDLLTGslehhhhhh | 0.12 |
| NF1-10 | mGSADQIIEDAQRQSPSVQVRKVSTPDEMDQAVRDVKNTGAQVVILVYTSSGELLIFAVTDPE<br>DARQIEEDAKKRNPSAKVVRLEGKPPDEIKKQMEKLLRGslehhhhhh | 1.3 |
| NF1-11 | mGTADDVMQDAKDQDPNAQVYKATTPDEIREAVKRIEKTGAQVVLIIYTSSGILILVAVTNPE<br>DADKILKEAKKRNPNVTLVRLEGKPPEDIRKQAENVWKGslehhhhhh | 0.65 |
| NF1-12 | mGDLDNMEKRAKELSPSVQIYRVTTPDEMEEAVQRVKNTGAQVVILVYTSSGRLIIFAVTNP<br>EIADRIKEDAKRQNPSAQLVRLEGASPDQIRQQIEDLLRGslehhhhhh | 0.34 |
| NF1-13 | mGTVEEIKKDAQKLDPNAQVREVTTPDEIEEAVRQVKNYGAQVVLLFYTSSGKIVVVAVRSK<br>EVADRIAQQVKDRNPSVTVIRLEGASPEEIREQMERLWKGslehhhhhh | 0.022 |
| NF1-14 | mGDADKIMEQAKRQDPNAQVYKVTTPDEIEEAVRRIEKYGAQVVLIIYTSSGIVILVAVRDPS<br>QADQILKEAKKQNPSATFVRLEGVSPDDLRRQVEDVWRGslehhhhhh | 0.94 |
| NF1-15 | mGDLENIIEKAKRQNPSAQVYEATTPDELDEVAERVQRTGAQVVILVYTSSGKIIVFAVTNPE<br>DAKRIVDEAKNQNPSAKVVRLEGASEDDMKEQMRRLWKGslehhhhhh | 2.1 |
| NF1-16 | mGDIENIIKDAKKQSSSVQVEKVTTPDEAEEVVRRVEKTGAQVVVLVYTSSGLVIVFAITNPEI<br>AKRIVQRAKEQNPSATVKRLEGVSPDEIQEEIERLLKGslehhhhhh | 0.17 |

**Supplementary Table 1 | Designed sequences of the series of NF1.**
"E-value" column shows the smallest E-value obtained from a PSI-BLAST search against nr database.

| ID | sequence | E-value |
|---|---|---|
| NF2-01 | mgSEIRLESSDGQDKTYTATSDDELKEILERAVKEGIKRIEIRGASERMLRTSEEIARRAGIEW RKNGslehhhhhh | 2.8 |
| NF2-02 | mgTEIELESKNGQREHYTATSEDEARKIIEKAVRRGIKRIELRGASEQLIRDMQEIAKQIGLQY RTDGslehhhhhh | 4.2 |
| NF2-03 | mgTKIHAEGPNGETRTYTATSEEEAEKIIRELVKRGIQRIELQGASEDLLRKMEEIARRAGIEY RTDGslehhhhhh | 0.015 |
| NF2-04 | mgKTIEAESSDGETRTYTATSEEEAERIIRKLQKEGIQRIRLQGVSEDLRKRLEELARKIGLQW RYRKgslehhhhhh | 0.16 |

**Supplementary Table 2 | Designed sequences of the series of NF2.**
"E-value" column shows the smallest E-value obtained from a PSI-BLAST search against nr database.

| ID | sequence | E-value |
|---|---|---|
| NF3-01 | mGDDDSLRKRLEEDAKKSGKRVEFRRYNDPKRIEEELRRARKDGETLVILVGGVIVIVSNDE KLVREIKKNLQKERPDKETISVTTEEDIKRALKRIKEGswslehhhhhh | 0.32 |
| NF3-02 | mGSSEKIRKKLEELAKRTGKRVQFREYNDTEQVRKTLEEAQRRGETLVVLSRGTVIIVSTNEE LIRRIEELVKQSNPNLETYEATTDEDIERILRELDKGslehhhhhh | 0.34 |
| NF3-03 | mGSDEEIRKKLEELAKRKGKDLQLRRYNDPNEVEKSIREALKKGRTLIIIINGVFVVVSTDEDL IREIKRLIKESNPNKKTLDVTTEEDLEEVLRRIKKGswslehhhhhh | 0.001 |
| NF3-04 | mGDSERLERRIRERAKKTGKDLQFREYDSPDKVRESLRKAQEKGRTLIVLVRGTIIVVSPDPE LARQIVEDLQKERPDLTTLEATTEDDIRKQLKRLREGswslehhhhhh | 0.52 |

**Supplementary Table 3 | Designed sequences of the series of NF3.**
"E-value" column shows the smallest E-value obtained from a PSI-BLAST search against nr database.

| ID | sequence | E-value |
|---|---|---|
| NF4-01 | mGSEEIYRLIEKIWRDIKNENPNARILIFIVFTSDGKIEVIIVIIDDDEELLRRIEETAKKRVPKV EIRRDRNEERAQKKIEEELKKRNPNATLYTVTSLDELEEILKKLTKQEGslehhhhhh | 4.2 |
| NF4-02 | mGSRKVYELVRKVWEAIKEENPNAKILIFLLFTSDGTIQIIIVIIDSSEETLRRIEEQIRKRVPNV RIERSKNEEEARKEIEKELKDRDPNATLYEVTTKEELKKILEKLERQEGslehhhhhh | 0.14 |
| NF4-03 | mGSERIYKIIKEIWETIKKENPKVKILIFILLTSDGTIEIIIVVISSNREEAQRIVEELQKRYPEVEI RRSENEEQASREIKQELKDRNPTATIYEVTSPEELTKILEKLKKQEGslehhhhhh | 0.88 |
| NF4-04 | mGSEEIRELVRKIYETVRKENPNVKILIFIIFTSDGTIKVIIVIIADDPNDAKRIVKKIQERFPKLT IKQSRNEEEAEKRIQKELEERNPNAEIQVVRSEDELKEILDKLDEKKGswslehhhhhh | 0.52 |
| NF4-05 | mGEDELKEQITRVWRTIKKENPGVKILVFILFSSNGEIQVIIVIISDNEDELRELEERAKRRVPK VEIRRDRDEQKASDKIREELKRRDPNAEIFEVTSEEELKKIIEELKRQKGslehhhhhh | 6.6 |
| NF4-06 | mGTDKVEELVRKIYESVEKENPNAKILIFLVYTSDGKILIIIVIISSSEETAKKIVEELKRRFPEV EIRQDRDEESAKKKIEEELRRRDPNATIYKVTSKDELEKILEKLRRDEGslehhhhhh | 0.18 |

**Supplementary Table 4 | Designed sequences of the series of NF4.**
"E-value" column shows the smallest E-value obtained from a PSI-BLAST search against nr database.

| ID | sequence | E-value |
|---|---|---|
| NF5-01 | mGEEERVKREAKRIEDEDPNRKILIYIDSNGEIEIKEVTSPEDVRKILEKLGVSEDLLREIER AVKNGEYDLFFIVKTEESTRRAEEIRERLGKPVRIETGslehhhhhh | 0.44 |
| NF5-02 | mGEEEKILRKADKIIKEDPNRKIIIIINPDGKIELREVTSEEDVREILERMGVPPDLLKEIERA VRNGEYDLFFIVTTEESERRARKLKKQMGKPVLIETGswslehhhhhh | 0.039 |
| NF5-03 | mGEDDEILQRAKDILKEDPNRKILIILNPDGKIELYEVTSEEDIKRIAKKAGISEELLRRILQS FRDGQYDLFFIAKTEDDERRARELKERMGKPVEILRGslehhhhhh | 0.36 |
| NF5-04 | mGDKEKVEETFRKIEEEDPNRKIIIIINPDGKIEIRTVTSPEDLERIFRKMGISEDLLRTAKE SLREGQYDLFFITKTEESRRTAEELKKRLGKPVLILTGswslehhhhhh | 0.2 |
| NF5-05 | mGNDEKVKETARKLLEKNPKVKIIIILNPDGEIRVKTVTSPDDLEQIARESGLPPDLLEQAL RDLKNGQYDLFVFAKTEEDERRARQLKEDMGKPVLILRGswslehhhhhh | 2.3 |
| NF5-06 | mGNEDEARERIRKIEDEDPNAKIIIYLTPDGELRVQKVTSEDDVEEFLKKMGVPPELLKRI LESLRNGEYDLFFIVRTEESEKRARKLKERLGKPVEILRGslehhhhhh | 1.1 |

**Supplementary Table 5 | Designed sequences of the series of NF5.**
"E-value" column shows the smallest E-value obtained from a PSI-BLAST search against nr database.

| ID | sequence | E-value |
|---|---|---|
| NF6-01 | mGELYTVDSPDEVRRIAKELGLSEEQLRRIEKEFRRAEKKGKTVLVYIDSNGEVRIREVTSE DELRELLQRLGVDPEIIERIERKFNNGEIKLVIIKGslehhhhhh | 0.11 |
| NF6-02 | mGKLYEVDSPDSVEKIARELGLSEEQLRRIQKEFERAERKGKLVIVYLTSDGKVEIREVTSE EELEKILKKLGVDEEIIRRIKRLRKEGQIKLVIIEGslehhhhhh | 0.002 |
| NF6-03 | mGELRTVDSPDSVREIIKRLGLSEEQLRRIEKKFKELEKKGRTLLVYIDSNGNVELRTVTSE DELERILRELGVDEEILRRVKELFREGQVKLVIIIGswslehhhhhh | 0.15 |
| NF6-04 | mGELRTVTSEDEVEKIARQLGLSEEQLRRILKKFREVEKRGKLLIVYLTSDGEVRIQEVTTK DELSRILKELGVDPEIRERIRKEFEEGQVKLVFIKGswslehhhhhh | 0.1 |
| NF6-05 | mGELRTVTTKEDVRRFAKKTGLSEDQLRKIEERFEEAEKKGKTIIVYITSDGKVEIQEVDTE DELKRILNELGVDPDIKEKIRKKFENGEVKLVFILGswslehhhhhh | 0.24 |
| NF6-06 | mGKLYRVDSPDSVRKIAKELGLSEEQLRRIEKSFKEVEKKGKTLIVWLDSNGNVEEQTVDT KDELSRILNELGVDPEIKERIRKLFENGEVKLVIIEGslehhhhhh | 0.26 |

**Supplementary Table 6 | Designed sequences of the series of NF6.**
"E-value" column shows the smallest E-value obtained from a PSI-BLAST search against nr database.

| ID | sequence | E-value |
|---|---|---|
| NF7-01 | mGLVQRFDVDENSEQVERLIRIAGLDEDKFEKAEIIVIVIVKTEEKLKRLAQRVKDLGADIILE INMDENSETVKRLAKEAGIPPDELRRAEILILVLVKTEEKAQRISEQIKRQGswslehhhhhh | 0.15 |
| NF7-02 | mGETTQFDVDENSEKVKRLIRKAGLSEEELKKADIIVIVISRNPEELKRLEEIVRNLGADRIIK LNVDENPEQVRQFAEEAGIPPEKLKRIDYLVVIISKTKEEAKELAERIKRQGswslehhhhhh | 0.27 |
| NF7-03 | mGTTRQYDVDENPETVEKLIRIAGIPKEELDRADIIVFILSRSEEKARRLKKKIQSLGADRIQII NVDENSEEVKRFAKTAGISEEELRKSEYLIIVISKTKDEAKRLSEEIKKQGslehhhhhh | 0.48 |
| NF7-04 | mGQIQYFNVDENPEQVRKLIEQAGLDPDELREAEVIIIIISRTPEQLEKLSRQVKELGADRLLE FNVDENPEQASKLAKTAGISEKQLREADYIILILVRDEKKAKKFADSLRKKGslehhhhhh | 0.51 |
| NF7-05 | mGDVQRFDVDENSEQVERLARQAGLSEDELRKAKIIIILISRTEKKLRELEEQVKRLGADRWI LLNVDENPEQVEKLIRTAGISEDEFKRSEIIILILSKTKDEAEELSRRLKKTGslehhhhhh | 2.3 |
| NF7-06 | mGTVRRYDVDENSDQVKNLIKIAGLDPEKLKRSEIIVIIVVKTPEKLKRLSQQVKDLGADIIIEI NVDENSEQVQRLADEAGIPPEDLKKAEYLIFVLSKTKDKAEQISRDLEKKGslehhhhhh | 0.012 |

**Supplementary Table 7 | Designed sequences of the series of NF7.**
"E-value" column shows the smallest E-value obtained from a PSI-BLAST search against nr database.

| ID | sequence | E-value |
|---|---|---|
| NF8-01 | mGTILIFLDKNKEQAEKLAKEVGVTEIYESDNLEELYREIKERIERENPNATILTVTDPNELKKIQDEGKVDRIILLIKGslehhhhhh | 1.3 |
| NF8-02 | mGTIVIFLVENEERARRIAKELGATEIYKSDNLEEAERQISKELKKENPNAEILTVTDPEEVRRRREEGQLDRLIVIIKGswslehhhhhh | 3.1 |
| NF8-03 | mGTYLIFLTDDKKTAEELARKLGVTEIYESDNLEELLRRIQEEIKKRNPNAEILTVTNPDEVRRIKEEGKVDRLILIIRGslehhhhhh | 0.57 |
| NF8-04 | mGTILIFFSKSEETAERIAKELGATEIYKSDNEEEALRRLSKEIKKRNPNAEIRYTTNEDEVRKRQKNGQIDRLIIVILGslehhhhhh | 1.4 |
| NF8-05 | mGTILIFFSKNPEEIERLAKELGATRIYESDNEEEAYRRLEEQIRRENPNATVLRVTSPSEIRQKKKEGKVDLLIFVLIGslehhhhhh | 0.62 |
| NF8-06 | mGTILIFFTKDEETLRKLAKEVGVTEIYESDNEDEAYKRLKEEIRKRNPNAEILRVTDENELRRLRKEGQVDQFILVLIGslehhhhhh | 0.22 |
| NF8-07 | mGTIILFFTTDPNEAKKVYKELGATRITRSDNEEEASRRLKEEIERRNPNAKIYRYTTPEEARRTAEKEGATEIIIILIGslehhhhhh | 2.0 |
| NF8-08 | mGTIVIIFVTDPKEAEDIYQQLGATRITTSDNEDEARKELSRELKEENPNAKIYSYTTEDEAERTAREQGVTRIIIVLVGslehhhhhh | 1.5 |
| NF8-09 | mGTIIIFFTTNPEEAEKVYKKLGATEITTSRNEDEARERLEEKIRRENPNAKIYSTTNPDDAEKTAKNQGATRIIIVLIGslehhhhhh | 0.13 |
| NF8-10 | mGTIIIIFSTNPEDVKTIYKKLGVTRLTRSDNEDEARKRLSEEIKRENPNAKIISTTTPDEAKREQEQQGATRIILVEIGswslehhhhhh | 0.57 |
| NF8-11 | mGTIIIIIVTDPNEAKTIYKELGATEIYESDNEEELLRRIEEQLRRENPNAKIYTTKNKEEAERTAREQGATRIIILLVGslehhhhhh | 1.1 |
| NF8-12 | mGTIIIIFVTDPNEAQRIYDELGATRIYKSRNEDEAREELRREIEEENPNATIISTKSEKEARKQQETQGATEIIIVLIGslehhhhhh | 2.5 |

**Supplementary Table 8 | Designed sequences of the series of NF8.**
"E-value" column shows the smallest E-value obtained from a PSI-BLAST search against nr database.

| Design ID | Expressed | Soluble | αβ-protein CD spectrum (20 °C) | Monomeric | Well-resolved HSQC |
|---|---|---|---|---|---|
| NF1-01 | Y | Y | Y | N | |
| NF1-02 | Y | Y | Y | Y | Y |
| NF1-03 | Y | Y | Y | N | |
| NF1-04 | Y | Y | Y | N | |
| NF1-05 | Y | N | | | |
| NF1-06 | Y | N | | | |
| NF1-07 | Y | Y | Y* | N | |
| NF1-08 | Y | Y | Y | Y | N |
| NF1-09 | Y | Y | Y* | N | |
| NF1-10 | Y | Y | Y | Y | Y |
| NF1-11 | Y | Y | Y* | N | |
| NF1-12 | Y | Y | Y | N | |
| NF1-13 | Y | Y | Y | Y | Y |
| NF1-14 | Y | Y | Y | Y | Y |
| NF1-15 | Y | Y | Y | Y | N |
| NF1-16 | Y | Y | Y* | N | |

**Supplementary Table 9 | Experimental summary of a series of designs for NF1.**

Each row corresponds to the results for each design. The columns give the results for each experimental characterization, of which the summaries are described in Extended Data Table 1. Each characterization was performed sequentially from the left to the right; well-behaved designs at a characterization (Y) are then evaluated by the next one and not well-behaved designs (N) end being evaluated.

* The CD spectrum was characteristic of αβ-proteins, but looked partially unfolded.

| Design ID | Expressed | Soluble | αβ-protein CD spectrum (20 °C) | Monomeric | Well-resolved HSQC |
|---|---|---|---|---|---|
| NF2-01 | Y | Y | Y | Y | N |
| NF2-02 | Y | Y | Y | Y | Y |
| NF2-03 | Y | Y | Y | Y | Y |
| NF2-04 | Y | Y | Y | Y | N |

**Supplementary Table 10 | Experimental summary of a series of designs for NF2.**

The summary was given in the same way as Supplementary Table 9.

| Design ID | Expressed | Soluble | αβ-protein CD spectrum (20 °C) | Monomeric | Well-resolved HSQC |
|---|---|---|---|---|---|
| NF3-01 | Y | Y | Y | Y | Y |
| NF3-02 | Y | N | | | |
| NF3-03 | Y | Y | Y | Y | Y |
| NF3-04 | Y | Y | Y | Y | † |

**Supplementary Table 11 | Experimental summary of a series of designs for NF3.**

The summary was given in the same way as Supplementary Table 9.

† The HSQC measurement was not conducted due to low concentration.

| Design ID | Expressed | Soluble | αβ-protein CD spectrum (20 °C) | Monomeric | Well-resolved HSQC |
|---|---|---|---|---|---|
| NF4-01 | Y | Y | Y | N | |
| NF4-02 | Y | Y | Y | Y | ‡ |
| NF4-03 | Y | Y | Y | Y | Y |
| NF4-04 | Y | Y | Y | Y | Y |
| NF4-05 | Y | Y | Y | Y | Y |
| NF4-06 | Y | Y | Y | Y | Y |

**Supplementary Table 12 | Experimental summary of a series of designs for NF4.**

The summary was given in the same way as Supplementary Table 9.

‡ The HSQC measurement was not conducted due to not small amount of dimeric state (the second peak of SEC-MALS).

| Design ID | Expressed | Soluble | αβ-protein CD spectrum (20 °C) | Monomeric | Well-resolved HSQC |
|---|---|---|---|---|---|
| NF5-01 | Y | Y | Y* | N | |
| NF5-02 | Y | Y | Y* | N | |
| NF5-03 | Y | Y | Y | Y | Y |
| NF5-04 | Y | Y | Y | N | |
| NF5-05 | Y | Y | Y | N | |
| NF5-06 | Y | Y | Y | Y | Y |

**Supplementary Table 13 | Experimental summary of a series of designs for NF5.**

The summary was given in the same way as Supplementary Table 9.

* The CD spectrum was characteristic of αβ-proteins, but looked partially unfolded.

| Design ID | Expressed | Soluble | αβ-protein CD spectrum (20 °C) | Monomeric | Well-resolved HSQC |
|---|---|---|---|---|---|
| NF6-01 | Y | Y | Y | Y | Y |
| NF6-02 | Y | Y | Y | Y | Y |
| NF6-03 | Y | Y | Y | N | |
| NF6-04 | Y | Y | Y | Y | Y |
| NF6-05 | Y | Y | Y | Y | Y |
| NF6-06 | Y | Y | Y | Y | Y |

**Supplementary Table 14 | Experimental summary of a series of designs for NF6.**

The summary was given in the same way as Supplementary Table 9.

| Design ID | Expressed | Soluble | αβ-protein CD spectrum (20 °C) | Monomeric | Well-resolved HSQC |
|---|---|---|---|---|---|
| NF7-01 | Y | Y | Y* | N | |
| NF7-02 | Y | Y | Y | Y | Y |
| NF7-03 | Y | Y | Y | Y | Y |
| NF7-04 | Y | Y | Y | Y | Y |
| NF7-05 | Y | Y | Y* | N | |
| NF7-06 | Y | Y | Y* | N | |

**Supplementary Table 15 | Experimental summary of a series of designs for NF7.**

The summary was given in the same way as Supplementary Table 9.

* The CD spectrum was characteristic of αβ-proteins, but looked partially unfolded.

| Design ID | Backbone type | Expressed | Soluble | αβ-protein CD spectrum (20 °C) | Monomeric | Well-resolved HSQC |
|---|---|---|---|---|---|---|
| NF8-01 | 1 | Y | Y | Y | Y | Y |
| NF8-02 | 1 | Y | Y | Y | Y | Y |
| NF8-03 | 1 | Y | Y | Y | N | |
| NF8-04 | 1 | Y | Y | Y | Y | Y |
| NF8-05 | 1 | Y | Y | Y | Y | Y |
| NF8-06 | 1 | Y | Y | Y | Y | Y |
| NF8-07 | 2 | Y | N | | | |
| NF8-08 | 2 | Y | Y | Y* | N | |
| NF8-09 | 2 | Y | Y | Y | N | |
| NF8-10 | 2 | Y | Y | Y | N | |
| NF8-11 | 2 | Y | Y | Y | Y | Y |
| NF8-12 | 2 | Y | Y | Y | N | |

**Supplementary Table 16 | Experimental summary of a series of designs for NF8.**

The summary was given in the same way as Supplementary Table 9. The backbone type 1 has GABA loop immediately before the last strand and 2 has GBA loop (see Extended Data Fig. 4 for details).

* The CD spectrum was characteristic of αβ-proteins, but looked partially unfolded.

#### References

1. M. Zweckstetter, A. Bax, Prediction of sterically induced alignment in a dilute liquid crystalline phase: Aid to protein structure determination by NMR. *Journal of the American Chemical Society* **122**, 3791-3792 (2000).
